# TNIP1 inhibits Mitophagy via interaction with FIP200 and TAX1BP1

**DOI:** 10.1101/2022.03.14.484269

**Authors:** François Le Guerroué, Achim Werner, Chunxin Wang, Richard Youle

## Abstract

Mitophagy is a form of selective autophagy that disposes of superfluous and potentially damage-inducing organelles in a tightly controlled manner. While the machinery involved in mitophagy induction is well known, the regulation of the components is less clear. For example, it is still unknown how the ULK1 complex is dissociated prior to closing of the autophagosome. Here, using Chemically Inducible Dimerization (CID) and mitophagy assays with the fluorescent probe mtKeima, we reveal that the ubiquitin binding domain-containing protein TNIP1 is able to induce mitophagy when ectopically targeted to mitochondria. Conversely, we demonstrate that TNIP1 knock down accelerates mitophagy rates, and that ectopic TNIP1 negatively regulates the rate of mitophagy. These functions of TNIP1 depend on a previously unrecognized, evolutionarily conserved LIR motif as well as its AHD3 domain, which are required for binding to the ULK1 complex member FIP200 and the autophagy receptor TAX1BP1, respectively. Taken together, our findings identify a novel negative regulator of mitophagy that acts at the early steps of autophagosome biogenesis and provide a molecular rationale for how the ULK1 complex might be dissociated from the closing autophagosome.

## Intro

Macroautophagy, one of the main cellular degradation pathways, sequesters cytosolic components inside a double membrane structure called an autophagosome, that matures, closes and catabolizes its content by fusing to lysosomes. The molecular mechanisms of the formation and maturation processes are well described and revolve around a set of proteins called the ATG conjugation system^1^. Initially described as a non-selective degradation process triggered when cells face starvation, more recent work shows autophagy can selectively eliminate certain proteins, protein aggregates and organelles^2,3^. Selective autophagy is the specific sequestration of cytosolic structures mediated via autophagy receptors. The main feature of these autophagy receptors is their capacity to bind to mATG8 (mammalian ATG8 proteins) as well as FIP200 and to ubiquitin in most cases^4^. One of the best characterized selective autophagy processes is the degradation of mitochondria via mitophagy^5^. The PINK1/Parkin axis of mitophagy participates in a feed-forward loop involving stabilization of the kinase PINK1, phosphorylation of resident ubiquitinated proteins at the surface of the mitochondria prior to recruitment of the E3 ligase Parkin, followed by activation of Parkin by its phosphorylation by PINK1. Autophagy receptors such as NDP52 or OPTN are subsequently recruited to ubiquitinated mitochondria via their ubiquitin-binding domains, allowing the autophagy machinery to assemble and degrade the mitochondria in a wholesale fashion^6,7^.

TNIP1 (TNFAIP3-interacting protein 1), also called ABIN1 (A20-binding inhibitor of NF-κB activation 1) participates in the NF-κB pathway, where it negatively regulates NF-κB activation, acting as a security check to maintain immune homeostasis^8,9^. Structurally, TNIP1 possesses a Ub-binding domain (UBAN domain – Ubiquitin binding of ABIN and NEMO) common among ABIN proteins, 3 Abin homology domains (AHD) and a NEMO binding domain (NBD)^8,10^. Although the AHD3 domain function is currently not described, AHD1 mediates binding with the ubiquitin editing enzyme Tumor necrosis factor alpha-induced protein 3 (TNFAIP3, A20) and AHD4 mediates the interaction with OPTN^11^. TNIP1 also interacts with TAX1BP1, together with A20 to restrict antiviral signaling^12^. Being involved in immune responses, dysregulation of TNIP1 was shown to be implicated in various human diseases in a number of genome-wide associated studies (GWAS)^13^. TNIP1 single nucleotide polymorphisms (SNP) have been strongly associated with autoimmune diseases^14^. Furthermore, cross ethnic genetic studies identified the *GPX3-TNIP1* locus to associate with amyotrophic lateral sclerosis (ALS)^15^. A later study concluded that this locus was less likely to contribute to ALS risk^16^. In addition, a recent GWAS study identified TNIP1 at the locus as a newly identified risk allele for Alzheimer’s disease (AD)^17^.

To date, a role of TNIP1 in regulating mitophagy has not been described. Here we define the implication of TNIP1 as an LC3-interacting protein, that is able to trigger mitophagy when synthetically targeted to mitochondria, and that normally negatively inhibits early stages of mitophagy. Usp30 also inhibits mitophagy and is a promising drug targtet to increase mitochondrial quality control^18^. Furthermore, we show that physical interaction of TNIP1 with the ULK1 complex member FIP200, as well as with TAX1BP1, drives mitophagy inhibition. Our study suggests that the autophagy protein TAX1BP1 and FIP200 are not only involved in promoting mitophagy, but are also crucial targets for autophagy inhibition.

## Results

### Ectopic localization of TNIP1 to the mitochondria induces mitophagy

TNIP1 possesses a UBAN domain similar to that in OPTN^19,20^ and was suggested to be an autophagy substrate^21^. In addition, TNIP1 was previously shown to form a complex with TAX1BP1 in a non-autophagy context^12^. Thus, we investigated whether TNIP1 is involved in mitophagy by interaction with TAX1BP1. However, HeLa cells express TNIP1 and knockout of all five mitophagy receptors (OPTN, p62, TAX1BP1, NBR1 and NDP52) completely blocks mitophagy, indicating that TNIP1 is unlikely to act alone in mitophagy induction or its function is fully dependent on those receptors^22^. Therefore, we took advantage of the chemically induced dimerization (CID) assay in cells expressing mtKeima^22^ (**Figure 1A**). We fused the protein FRB to the C-terminus tail of the outer mitochondrial protein FIS1 and TNIP1 to the soluble protein FKBP. Adding the small molecule RAPALOG induces dimerization of FRB and FKBP, artificially localizing TNIP1 to the mitochondria. Interestingly, localizing TNIP1 to mitochondria for 24 Hrs induced robust mitophagy (**Figure 1B**). TNIP1 with the UBAN loss of function mutation (D472N) was able to trigger mitophagy to levels similar to WT TNIP1, indicating that the ubiquitin binding domain of TNIP1 is not necessary for inducing mitophagy when TNIP1 is artificially targeted to mitochondria.

**Fig. 1.**
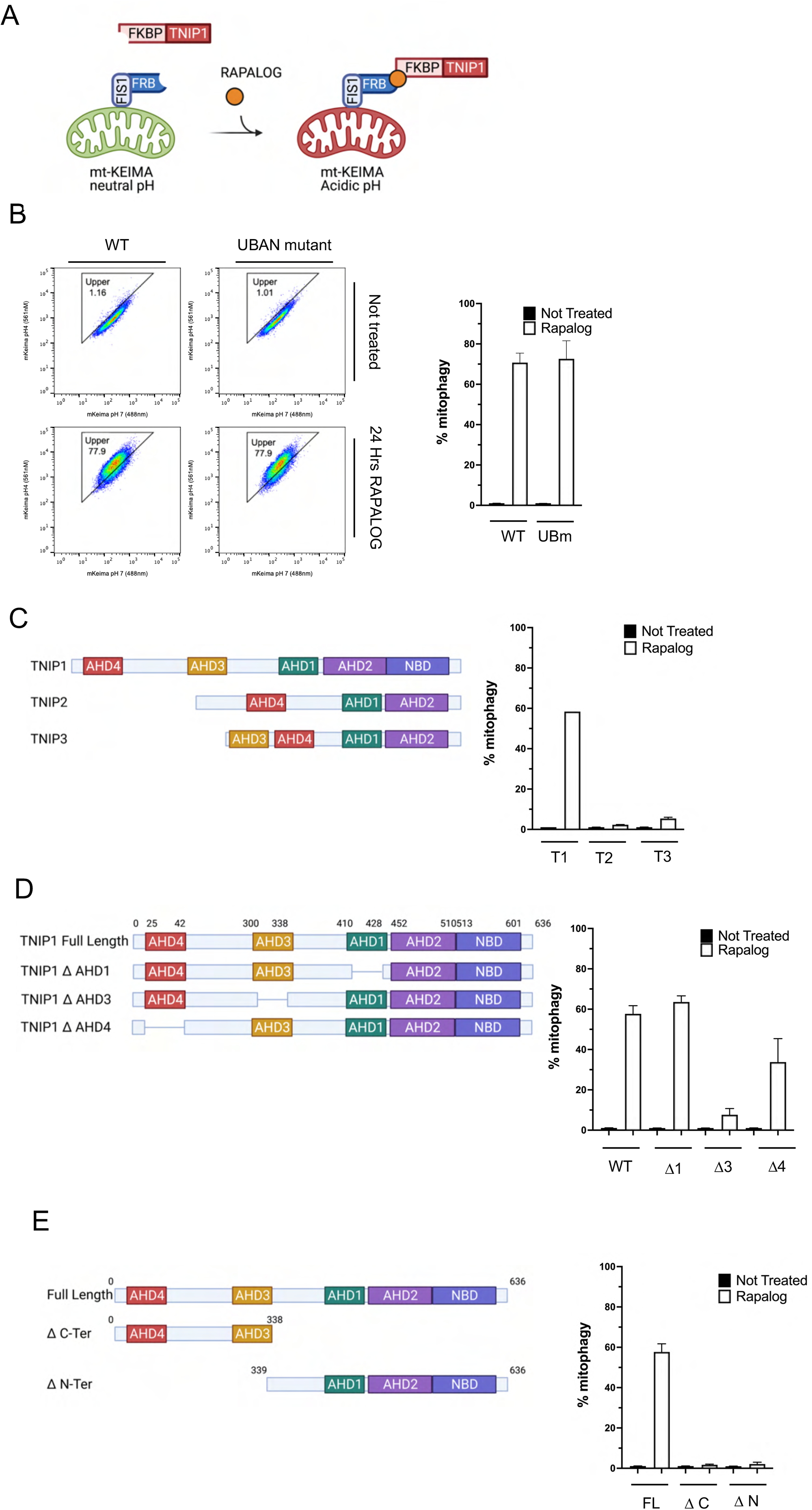
Ectopic localization of TNIP1 to the mitochondria induces mitophagy. (A) schematization of CID experiment, made with the help of BioRender. (B) Left, cells were treated with RAPALOG for 24 Hrs and subjected to FACS acquisition. FACS plot of mtKeima ratio (561/488) of HeLa cells stably expressing mito-Keima, FRB-FIS1 and FKBP-GFP-TNIP1 WT or UBAN mutant (D472N). Right, quantification of mtKeima ratiometric analysis of FACS data from 3 independent replicates. WT: Wild type. UBm: UBAN mutant (D472N). (C) Left, schematic representation of the TNIP family TNIP1, TNIP2 and TNIP3, made with the help of BioRender. NBD, Nemo Binding Domain. AHD2 is also named UBAN domain. Right, cells were treated with RAPALOG for 24 Hrs and subjected to FACS acquisition. Quantification of mtKeima ratiometric analysis (561/488) of HeLa cells stably expressing mito-Keima, FRB-FIS1 and FKBP-GFP-TNIP1 or FKBP-GFP-TNIP2 or FKBP-GFP-TNIP3 of FACS data from 3 independent replicates. T1: TNIP1. T2: TNIP2. T3: TNIP3. (D) Left, schematic representation of TNIP1 and the AHD mutant constructs, made with the help of BioRender. Right, cells were treated with RAPALOG for 24 Hrs and subjected to FACS acquisition. Quantification of mtKeima ratiometric analysis (561/488) of HeLa cells stably expressing mito-Keima, FRB-FIS1 and FKBP-GFP-TNIP1 WT or FKBP-GFP-TNIP1 mutants ΔAHD1, ΔAHD3 and ΔAHD4 of FACS data from 3 independent replicates. WT: Wild type. Δ1: ΔAHD1. Δ3: ΔAHD3 andΔ4: ΔAHD4. (E) Left, schematic representation of full length TNIP1 or the ΔC-terminal and N-terminal part of TNIP1, made with the help of BioRender. Right, cells were treated with RAPALOG for 24 Hrs and subjected to FACS acquisition. Right, quantification of mtKeima ratiometric analysis (561/488) of HeLa cells stably expressing mito-Keima, FRB-FIS1 and FKBP-GFP-TNIP1 full length or FKBP-GFP-TNIP1 ΔC-terminal or N-terminal of FACS data from 3 independent replicates. FL: full length. ΔC: ΔC-terminal. N: ΔN-terminal.

Mammalian cells possess 3 TNIP homologs, TNIP1, TNIP2 and TNIP3, sharing certain ABIN homology domains (AHDs) that characterize them (**Figure 1C**). The AHD1 domain has been shown to be essential for interacting with A20/TNFAIP3, a central gatekeeper in inflammation and immunity. CID and FACS analysis with TNIP2 and TNIP3 revealed that they were not able to induce mitophagy as substantially as TNIP1, indicating the singular role of TNIP1 in mitophagy. To further characterize the important domains of TNIP1, we carried out CID with versions of TNIP1 missing individual AHD domains. AHD1 deletion had no effect on mitophagy, suggesting that binding to A20 is not necessary for its role in mitophagy. TNIP1 with the AHD3 deletion showed a strong reduction in its ability to induce mitophagy, while deletion of AHD4 domain in TNIP1 showed moderate inhibitory effect on mitophagy (**Figure 1D**). Both N-terminal and C-terminal fragments of TNIP1 (**Figure 1E**) proved incapable of inducing mitophagy, indicating that bipartite domains are involved in mitophagy and AHD3 domain is essential.

To determine the molecular mechanisms of TNIP1 in mitophagy, we carried CID-induced mitophagy in different cellular backgrounds: FIP200 KO, WIPI2 KO, A20 KO and pentaKO^22^ cells (NDP52, OPTN, TAX1BP1, SQSTM1 and NBR1). PentaKO (5KO) cells were previously described to be deficient in mitophagy, owing to the dependence on recruitment of autophagy receptors to recruit the autophagy machinery. To our surprise, TNIP1 was able to mediate mitophagy in 5KO cells at a level similar to that in WT cells (**supp Figure 1A**). This implies that TNIP1 is able to recruit an essential effector of the autophagy machinery on its own and does not depend on further recruitment of autophagy receptors previously deemed critical for mitophagy response, including TAX1BP1. In FIP200 KO and WIPI2 KO cells, no mitophagy was observed (**supp Figure 1A**), consistent with their essential roles in general autophagy machineries. Because of the tight role TNIP1 plays with A20 in the NF-κB pathway, we explored mitophagy activation in an A20 KO background (**supp Figure 1A**). Consistent with the results seen with the AHD1 deletion mutant, A20 KO did not alter the mitophagy response, supporting the idea that TNIP1’s role in CID induced mitophagy does not depend on A20, and revealing a new activity of TNIP1 distinct from prior work on its role in inflammation. Mitophagy independent of autophagy receptors (5KO) indicates that TNIP1 likely acts downstream of or in parallel with autophagy receptor recruitment and can recruit mATG8 and the autophagy machinery independent of the autophagy receptors, including TAX1BP1 previously reported to bind TNIP1.

To clarify the role of TNIP1 requirement in mitophagy, we conducted different CID experiments in TNIP1 KO cells. Unfortunately, owing to the size of FIP200, we could not produce a functioning FKBP-GFP-FIP200 construct. Consequently, we assessed whether other autophagy machinery proteins (i.e. WIPI2 and ATG16L1), as well as A20 were able to trigger mitophagy when placed on mitochondria in WT or TNIP1 KO cells. As previously reported, CID with ATG16L1 showed a strong mitophagy response in control cells^7^. TNIP1 KO did not impair that response, and even slightly enhanced this response (**supp Figure 1B**). Similarly, CID with WIPI2 displayed a robust mitophagy response and this response was stronger in TNIP1 KO cells (**supp Figure 1B**). Finally, CID with A20 displayed a weak mitophagy response that was abolished in TNIP1 KO cells (**supp Figure 1B**). This could be explained by a weak recruitment of TNIP1 mediated by A20 to the mitochondria, and the subsequent recruitment of the autophagy machinery. In light of these results, we conclude that TNIP1 can induce mitophagy independently of autophagy receptors, but is dependent on early autophagy machinery and is dispensable for these autophagy machinery proteins to induce mitophagy.

### TNIP1 is a negative regulator of mitophagy

The CID results showed that TNIP1 has the potential to trigger mitophagy and allowed us to identify the domains of TNIP1 required for this function. Since TNIP1 possesses a ubiquitin-binding domain, we speculated that it could act as an autophagy receptor working in concert with TAX1BP1. To test this hypothesis, we compared TNIP1 to known autophagy receptors in mitophagy flux assays. When subjected to mitophagy induced by mitochondrial depolarization, receptors are degraded in an autophagy-specific manner, together with their cargo. Blocking autophagosomal degradation with the V-ATPase inhibitor Bafilomycin A1 (BafA1) is a common way to estimate substrates specifically degraded in lysosomes. Combining the mitochondrial ATP synthase and complex III inhibitors Oligomycin A and AntimycinA1 (OA), respectively, is an established treatment for triggering mitophagy. Autophagy receptors are selectively degraded during mitophagy, as can be seen with TAX1BP1, NDP52 and OPTN (**Figure 2A**). Double treatment of cells with BafA1 and OA promotes mitophagy, but prevents lysosomal degradation of the encapsulated proteins (**Figure 2A**). However, similar to the ULK1 complex protein FIP200, TNIP1 levels were not influenced by Bafilomycin or OA treatment. Interestingly, TNIP1 was recently found to be an autophagy substrate in an autophagosome profiling content screening^21^, and thus would appear to be an autophagy substrate. Consequently, its apparent lack of degradation upon mitophagy or absence of accumulation upon Bafilomycin A1 treatment may be due to a fast turnover rate. Therefore, we performed OA treatments combined with cycloheximide (CHX) to investigate TNIP1 degradation (**supp Figure 2A**). Remarkably, TNIP1 seems to be a long-lived protein, as no degradation of TNIP1 was seen after 4 hours of CHX treatment, indicating that TNIP1 does not seem to be rapidly turned over. Interestingly, under steady state condition, FIP200 is degraded rapidly but counterintuitively is stabilized during mitophagy induction. Consequently, TNIP1 is very likely not degraded *en masse* via autophagy, as is the case for FIP200. Immunocytochemistry allowed observation of TNIP1 localization under steady state condition as well as upon mitophagy. To trigger mitophagy, HeLa cells stably expressing HA-Parkin were treated for 4Hrs with OA (**Figure 2B+supp Figure 2B**). Under steady state condition, TNIP1 appeared as puncta, in the vicinity of the mitochondria, with some colocalization events at the periphery of mitochondria, in a manner reminiscent of autophagy receptors. However, upon OA treatment, TNIP1 was found to coalesce in the perinuclear region and not to substantially relocate to mitochondria, contrary to what is seen with the autophagy receptor TAX1BP1, prompting us to reconsider the role of TNIP1 as an autophay receptor.

**Fig. 2.**
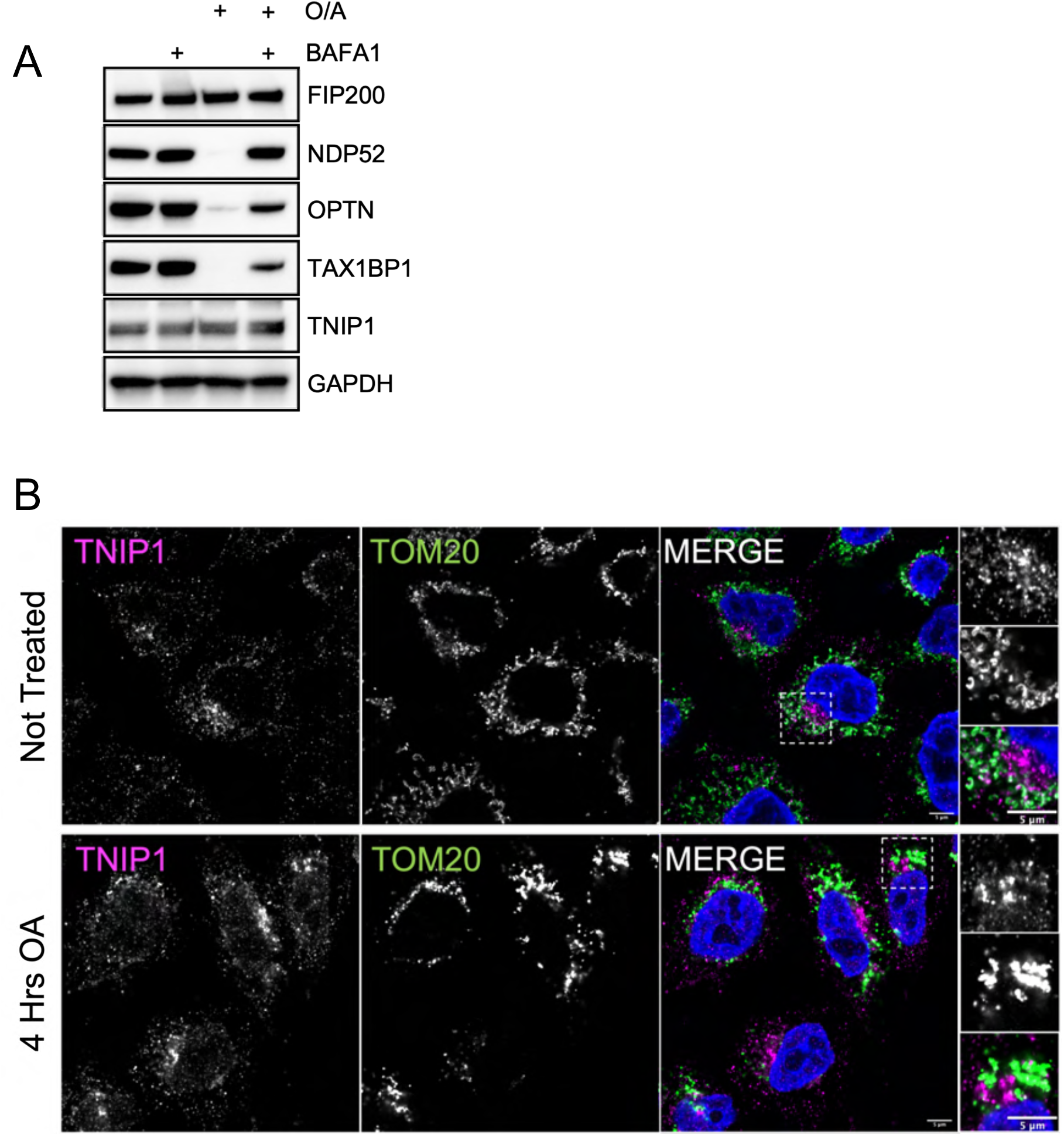
TNIP1 is not an autophagy substrate. (A) HeLa cells stably expressing HA-Parkin were treated for 4 Hrs with Oligomycin and Antimycin (O/A) and/or Bafylomycin A1 and subjected to immunoblot analysis. (B) HeLa cells stably expressing HA-Parkin were treated for 4 Hrs with Oligomycin and Antimycin (O/A) and stained for the mitochondrial protein TOM20 and endogenous TNIP1 before immunofluorescence acquisition on a confocal microscope. Airyscan representative images. Scale bar: 5μm.

Because TNIP1 was able to induce mitophagy when artificially placed onto mitochondria, we next asked if TNIP1 KO cells show any alteration in PINK1/Parkin mediated mitophagy. HeLa cells expressing mtKeima and HA-Parkin were treated for 6Hrs with OA and analyzed by FACS. This treatment induced a strong mitophagy response in WT cells and showed the same response in TNIP1 KO cells (**Figure 3A**). In contrast, reexpression of TNIP1 in TNIP1 KO cells using the same construct used for the CID experiments (FKBP-GFP-TNIP1) showed a substantial defect in mitophagy. These results indicate that TNIP1 may be a negative regulator of mitophagy and prompting us to investigate earlier time points in mitophagy. Indeed, at 2Hrs of OA treatment, when mitophagy was lower in WT cells, we saw a stronger mitophagy induction in TNIP1 KO cells. A response that could be reversed by re-expression of TNIP1 (**Figure 3B**). To further validate TNIP1’s role in inhibiting mitophagy, we overexpressed TNIP1 in HeLa cells and monitored mitophagy using mtKeima and FACS. Similar to results seen with rescue experiments in the TNIP1 KO background, over-expression of TNIP1 in WT HeLa cells induced a robust inhibition of mitophagy (**Figure 3C**). Overexpression of AHD domain deletion mutants in WT cells was then used to compare to the full length TNIP1 and assess their abilities to inhibit mitophagy. While the ΔAHD1 construct showed a mitophagy response similar to WT TNIP1, in accordance with a lack of function in previous CID experiments, ΔAHD3 displayed higher mitophagy, while ΔAHD4 showed an intermediate response compared to ΔAHD3 and WT TNIP1. These results are consistent with the results obtained with CID, where we identified the AHD3 domain as being important for TNIP1 function.

**Fig. 3.**
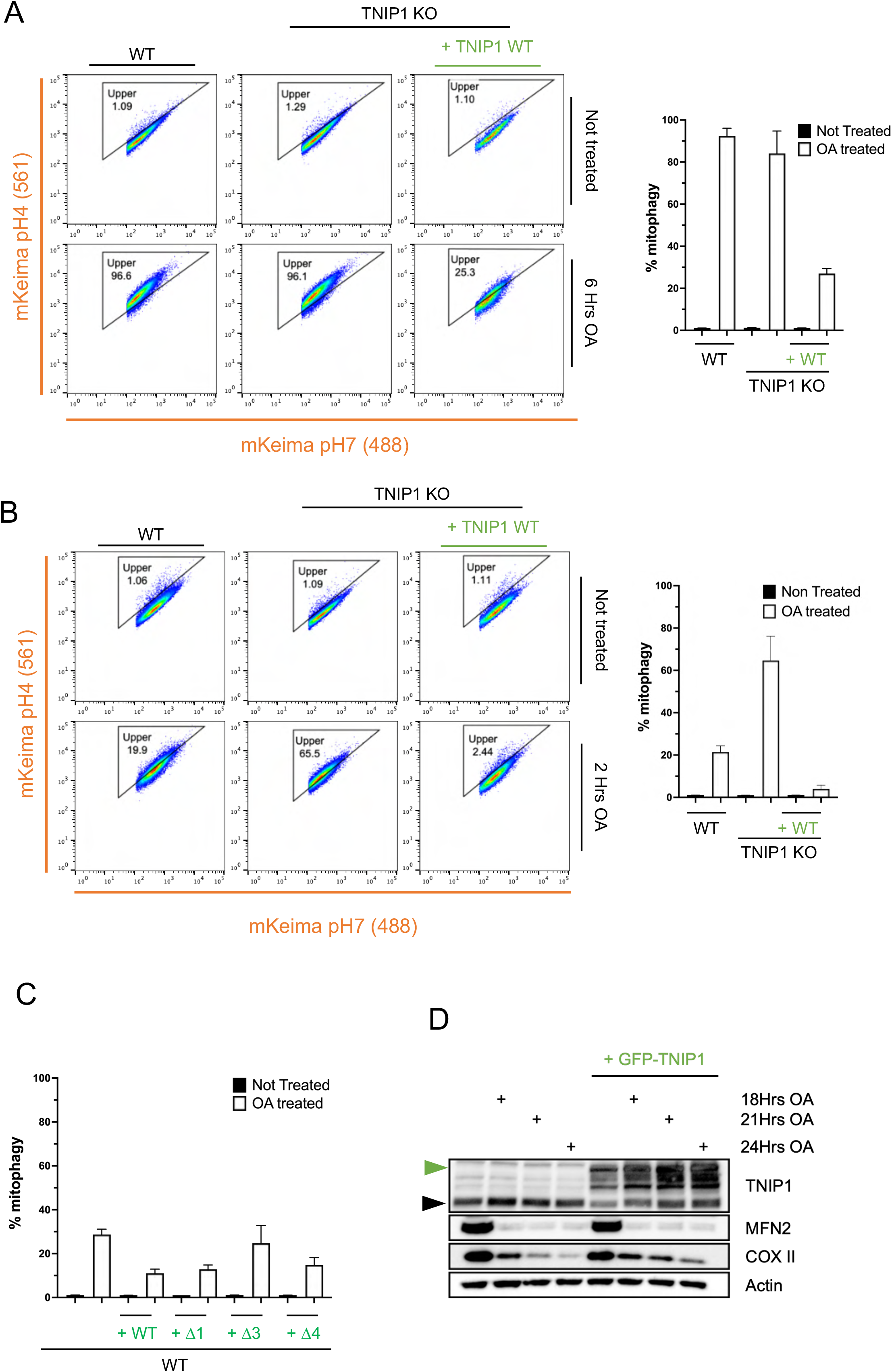
TNIP1 is a negative regulator of autophagy. (A) Left, cells were treated with Oligomycin and Antimycin (O/A) for 6 Hrs and subjected to FACS acquisition. FACS plot of mtKeima ratio (561/488) of HeLa cells stably expressing mito-Keima, HA-Parkin and FKBP-GFP-TNIP1 in wild type or TNIP1 KO cells. Right, quantification of mtKeima ratiometric analysis of FACS data from 3 replicates. WT: Wild type. + WT: + FKBP-GFP-TNIP1. These cells do not express the FRB-FIS1 construct and thus are not able to perform CID experiments. (B) Left, cells were treated with Oligomycin and Antimycin (O/A) for 2 Hrs and subjected to FACS acquisition. FACS plot of mtKeima ratio (561/488) of HeLa cells stably expressing mito-Keima, HA-Parkin and FKBP-GFP-TNIP1 in wild type or TNIP1 KO cells. Right, quantification of mtKeima ratiometric analysis of FACS data from 3 independent replicates. WT: Wild type. + WT: + FKBP-GFP-TNIP1. These cells do not express the FRB-FIS1 construct and thus are not able to perform CID experiments. (C) Cells were treated with Oligomycin and Antimycin (O/A) for 3 Hrs and subjected to FACS acquisition. Quantification of mtKeima ratiometric analysis (561/488) of HeLa cells stably expressing mito-Keima, HA-Parkin and FKBP-GFP-TNIP1 wild type, ΔAHD1, ΔAHD3 or ΔAHD4 constructs in wild type cells of FACS data from 3 independent replicates. + WT: + FKBP-GFP-TNIP1 WT, + Δ1: + FKBP-GFP-TNIP1 ΔAHD1, + Δ3: + FKBP-GFP-TNIP1 ΔAHD3, + Δ4: + FKBP-GFP-TNIP1 ΔAHD4. These cells do not express the FRB-FIS1 construct and thus are not able to perform CID experiments. (D) HeLa cells stably expressing HA-Parkin and FKBP-GFP-TNIP1 WT were treated for 18 Hrs, 21 Hrs or 24 Hrs with Oligomycin and Antimycin (O/A) and subjected to immunoblot analysis. Black arrow: endogenous TNIP1. Green arrow: FKBP-GFP-TNIP1.

To further confirm the specificity of TNIP1 in inhibiting mitophagy, we performed the same over-expression experiments with TNIP2 and TNIP3 (**supp Figure 3A**). While TNIP2 overexpression had no effect, TNIP3 overexpression showed a mild inhibition of mitophagy, but much less than seen with TNIP1 over-expression. This may be explained by the presence of an AHD3 domain in TNIP3 but not in TNIP2.

Another rigorous mitophagy assay is to monitor the degradation of the mitochondrial matrix protein COXII upon longer OA treatment. In WT cells, COXII showed a robust degradation after 18Hrs, progressing further after 21Hrs or 24Hrs of mitochondrial depolarization (**Figure 3D**). When TNIP1 is overexpressed, we observed a block in COXII degradation starting at 21Hrs of treatment, confirming that TNIP1 overexpression reduces the rate of mitophagy. Ectopic expression of TNIP1 however did not seem to influence canonical autophagy upon starvation as degradation of NDP52 or TAX1BP1 was not blocked when TNIP1 was overexpressed (**supp Figure 3B**). This indicates that TNIP1 appears specific for mitophagy and potentially other selective autophagy pathways.

### Binding of TAX1BP1 to the AHD3 domain is required for TNIP1’s inhibition of mitophagy

To obtain mechanistic insights into the function of TNIP1 as a negative regulator of autophagy, we next performed an unbiased mass spectrometry (MS) screen to identify specific interactors of the AHD3 domain. We immuno-precipitated (IP) HA-TNIP1 full length, TNIP1 ΔAHD3 and TNIP1 ΔAHD4 and determined high confidence interaction partners (HCIPs) by MS followed by Comparative Proteomics Analysis Software Suite (compPASS) analysis as previously described^23,24^ (**Figure 4A, Supplementary Table**). While TNIP1 co-immunoprecipitated with many cell surface receptors and secreted proteins, Gene Ontology (GO) analysis revealed autophagy components as the only significantly enriched class of proteins. In particular, RAB11FIP5, the autophagy receptors TAX1BP1 and OPTN as well as the ULK1 complex component RB1CC1 (FIP200) were the most abundant HCIPs.

**Fig. 4.**
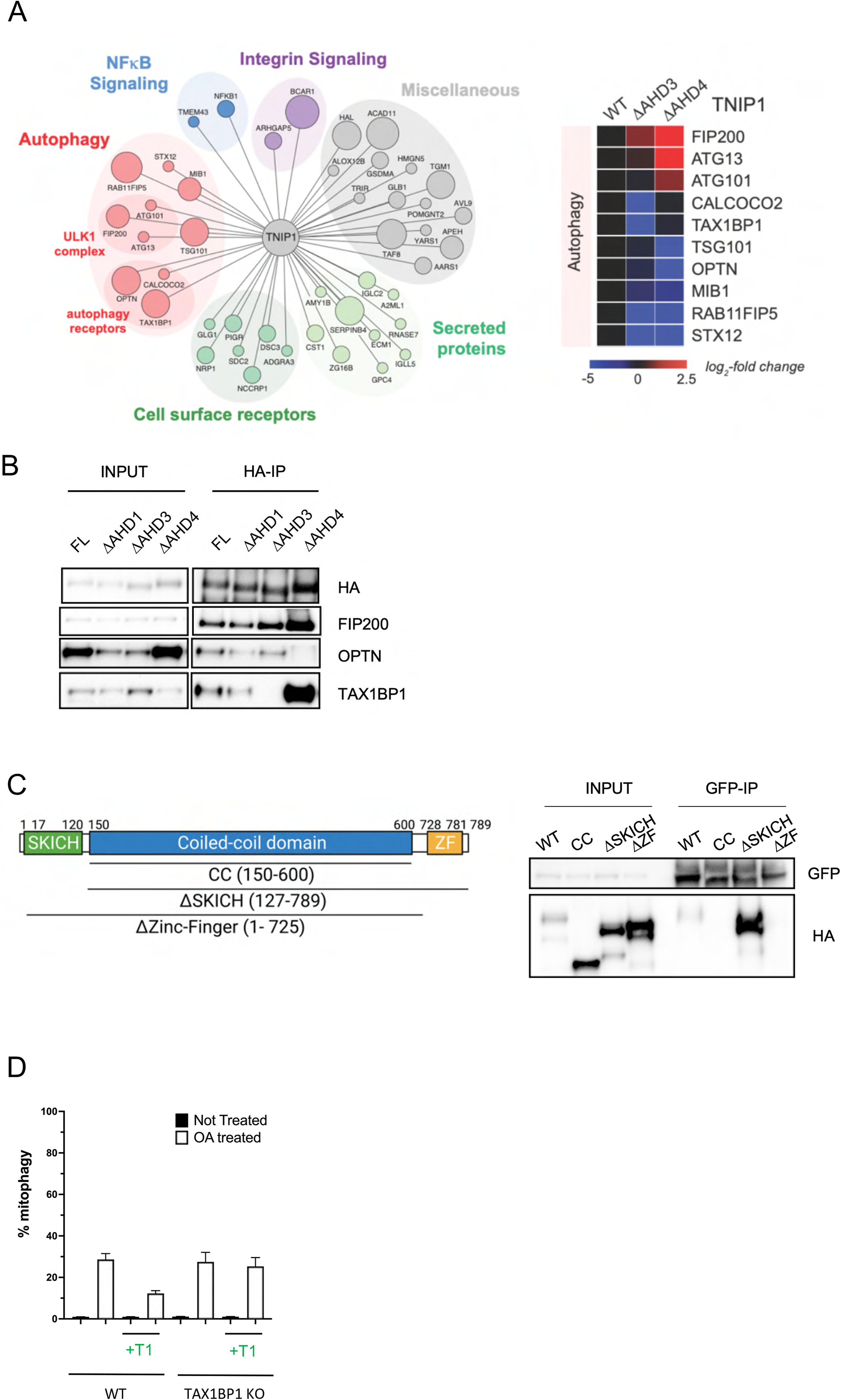
The interaction between the AHD3 domain of TNIP1 and the Zinc-Finger domain of TAX1BP1 is necessary for TNIP1’s mitophagy inhibition. (A) Left, High Confidence Candidate Interaction Proteins (HCIPs) network of HA-TNIP1 following an HA-pull down and mass spectrometry analysis. The size of the bait represent Higher Z score interaction. Right, Heat map of the interactors with wild type TNIP1 or ΔAHD3 and ΔAHD4 mutants. Gradient from blue to red reflects log2 fold change. (B) HEK293T cells stably expressing HA-TNIP1 full length or the ΔAHD1, ΔAHD3 and ΔAHD4 constructs were immunoprecipitated using magnetic HA beads and subjected to immunoblot analysis. (C) Top, schematic representation of full length TAX1BP1 and the regions encompassing the coiled-coil domain, ΔSKITCH and ΔZinc-Finger constructs, made with the help of BioRender. Bottom, HEK293T cells stably expressing GFP-TNIP1 full length and transiently expressing HA-TAX1BP1 full length, coiled-coil, ΔSKITCH and ΔZinc-Finger constructs were immunoprecipitated using magnetic GFP beads and subjected to immunoblot analysis. (D) Cells were treated with Oligomycin and Antimycin (O/A) for 3 Hrs and subjected to FACS acquisition. Quantification of mtKeima ratiometric analysis (561/488) of HeLa cells stably expressing mito-Keima, HA-Parkin and FKBP-GFP-TNIP1 in wild type or TAX1BP1 KO cells of FACS data from 3 independent replicates. WT: Wild type. + T1: + FKBP-GFP-TNIP1. These cells do not express the FRB-FIS1 construct and thus are not able to perform CID experiments.

Consistent with previous reports^11^, we observed that the TNIP1-OPTN interaction occurs via the ΔAHD4 domain by IP/MS and IP/immunoblotting (**Figure 4 A**,**B)**. Interestingly, we also confirmed binding of TNIP1 to TAX1BP1^12^, which we found to require the AHD3 domain, suggesting that TNIP1’s function in mitophagy might depend on TAX1BP1. To test this possibility, we further characterized the interaction between TNIP1 and TAX1BP1. We overexpressed HA-tagged TAX1BP1 full length, coiled-coil (CC), ΔSKICH or ΔZinc-Finger (ZF) constructs in 293T cells stably expressing GFP-TNIP1 followed by anti-GFP IPs (**Figure 4C**). these experiments revealed that TNIP1 is able to interact with full length TAX1BP1 and ΔSKICH TAX1BP1, but not with TAX1BP1 CC and ΔZF constructs. These results indicate that TNIP1 binds via the TAX1BP1 ubiquitin binding domain ZF. We next performed FACS experiments to assess the involvement of TAX1BP1 in TNIP1’s role in mitophagy inhibition (**Figure 4D**). No reduction in mitophagy was observed in TAX1BP1 KO cells compared to WT cells, in accordance to previously published data^22^. Furthermore, when we overexpressed TNIP1 in TAX1BP1 KO cells, we observed that TNIP1 did not inhibit mitophagy to levels similar to those in WT cells. These results indicate that TAX1BP1 is essential for TNIP1 inhibition of mitophagy. Overall, these results indicate that TNIP1 binds the ZF domain of TAX1BP1 via its AHD3 domain, and this binding is critical for its inhibitory action.

TANK-Binding Kinase 1 (TBK1) regulates several autophagy receptors in the autophagy pathway^7^. Furthermore, TBK1 was previously shown to be recruited to the TNIP1-TAX1BP1 complex^12^. We thus investigated whether TBK1 was involved in TNIP1 autophagy machinery recruitment. We performed CID experiments by targeting TNIP1 to mitochondria, coupled with immunocytochemistry and observed the recruitment of FIP200 and TAX1BP1 to the mitochondria in TBK1 KO cells (**supp Figure 4A**). Furthermore, we examined whether the absence of TBK1 prevented mitophagy inhibition seen with TNIP1 overexpression. (**supp Figure 4B**). Although we could see that TBK1 KO cells exhibited a marked reduction in mitophagy, ectopic expression of TNIP1 further reduced mitophagy. These combined results imply that TBK1 is likely not regulating TNIP1’s action. Perhaps these results are not surprising as TNIP1 does not seem to rely extensively on its UBAN domain for its function, while TBK1’s mode of action is principally to promote the association of UB chains to UB-binding domains through phosphorylation.

### Binding of FIP200 to an evolutionarily conserved LIR motif is required for TNIP1’s inhibition of mitophagy

The paradigm for selective autophagy initiation recently shifted, from the previous models that autophagy receptors role was to recruit mATG8 proteins, which in turn recruit the autophagy machinery, with the recent discovery that autophagy receptors can directly recruit the ULK1 complex for autophagosome formation^7,25,26^. Because the CID data using TNIP1 showed that mitophagy in 5KO cells still functioned (**supp Figure 1A**) and because of the co-IP Mass Spec data (**Fig. 4A**), we reasoned that TNIP1 may interact with components of the autophagy machineries other than TAX1BP1. While the main role of TNIP1 seems to be to inhibit mitophagy by preventing certain autophagy components to function, the synthetic placement of TNIP1 to the mitochondria in CID experiments seems to be enough to recruit these components and drive a robust mitophagy response. Other high confidence IPs in our MS data were FIP200 and other members of the ULK1 complex (ATG13 and ATG101). This binding was still observed with the ΔAHD3 and ΔAHD4 mutants. FIP200 binding to TNIP1 potentially explains how CID with TNIP1 induced mitophagy, as it was previously demonstrated that CID with a FIP200-binding peptide was sufficient to induce mitophagy^7^. We thus performed immunoprecipitation experiments to validate the binding of TNIP1 with FIP200 (**Figure 4B**). Full length TNIP1 was able to bind endogenous FIP200, as did ΔAHD1, ΔAHD3 and ΔAHD4 domain deletion mutants. SQSTM1/p62 was recently demonstrated to bind to FIP200 through its LIR motif ^25^. We thus investigated if TNIP1 contained an LIR motif that would also encompass a FIP200 binding region. In silico searches of LC3 interacting region (LIR)^27^ motif using the bioinformatic tool iLIR^28^ website revealed two potential canonical LIR domains in TNIP1 (**Figure 5A**). A Blast alignment with 4 other higher eukaryote TNIP1 proteins showed that LIR2 is conserved but not LIR1 (**supp Figure 5A**). In order to determine whether TNIP1 LIR2 is a functional LIR domain, we mutated LIR1 F83 and L86 to alanine and LIR2 F125 and V128 to alanine (**supp Figure 5B**). Lysates of HeLa cells transfected with TNIP1 WT, LIR1 mutant or LIR2 mutant were subjected to GST pull down experiments, an assay previously used to assess binding to mATG8 proteins (mammalian ATG8 i.e. LC3A, LC3B, LC3C, GABARAP, GABARAPL1 and GABARAPL2)^29,30^. As a positive control, SQSTM1/p62 was demonstrated to bind strongly to all mATG8 proteins (**supp Figure 5C**). TNIP1, on the other hand, was able to bind only some of the mATG8 proteins, with a preference for GABARAP, GABARAPL1 and GABARAPL2 (**Figure 5B**). LIR2 mutation abolished all binding to mATG8 proteins, while LIR1 mutations had no effect on binding, indicating that TNIP1 possess an ATG8 binding motif, characterized by the residues FEVV.

**Fig. 5.**
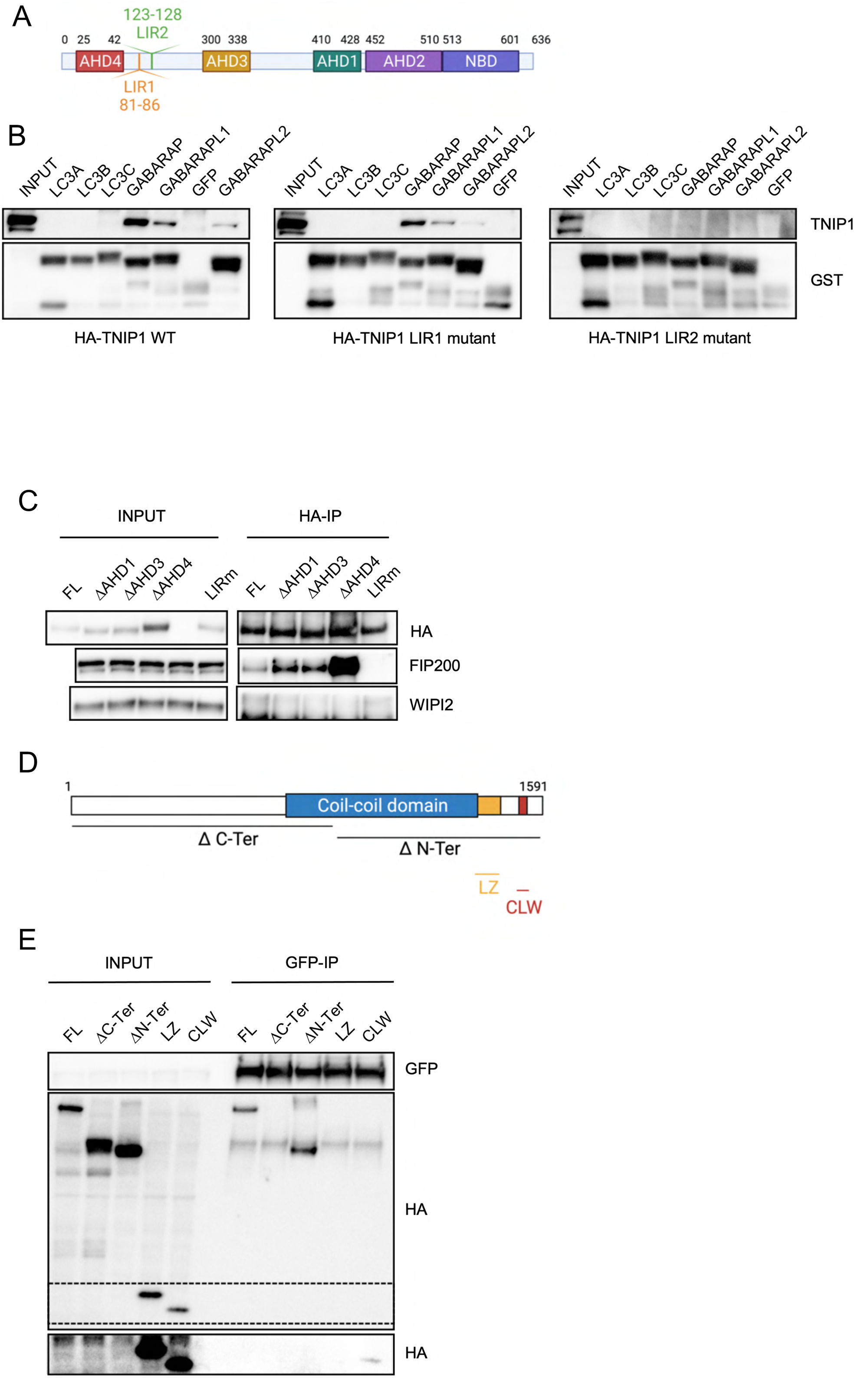
TNIP1 possess an LIR motif that is essential for it’s inhibitory role. (A) Schematic representation of TNIP1 and its 2 identified putative LIR motifs, made with the help of BioRender. (B) HeLa cells stably expressing HA-TNIP1 wild type, LIR1 mutant or LIR2 mutant were lysed and lysate was incubated with purified recombinant GST tagged mATG8 proteins of GFP. Precipitates were subjected to immunoblot analysis. Note that in the left IB, GFP and GABARAP lanes are inverted compared to the other two IB. (C) HEK293T cells stably expressing HA-TNIP1 full length or the ΔAHD1, ΔAHD3, ΔAHD4 and LIR mutant constructs were immunoprecipitated using magnetic HA beads and subjected to immunoblot analysis. (D) Schematic representation of full length FIP200 and the regions encompassing the N-terminal and C-terminal domains as well as the minimal leucine zipper (LZ) and Claw (CLW) domains, made with the help of BioRender. (E) HEK293T cells stably expressing GFP-TNIP1 full length and transiently expressing HA-FIP200 full length, N-terminal, C-terminal, LZ and CLW constructs were immunoprecipitated using magnetic GFP beads and subjected to immunoblot analysis. The zone in dotted lines was exposed longer to reveal the binding to the CLAW domain (lower part).

We next examined whether the TNIP1 LIR mutant construct was able to retain binding with FIP200 (**Figure 5C**). Reflecting the MS data that showed the ΔAHD4 mutant seemed to bind more strongly to FIP200 (**Figure 4A**), the ΔAHD4 mutant displayed a much stronger IP interaction with FIP200. Interestingly, we could not detect any FIP200 binding with the LIR mutant TNIP1. Reflecting the MS data where we did not detect a co-immunoprecipitation of TNIP1 with WIPI2, binding with WIPI2 was not detected. Similar to observations made with CID, where N-Terminal and C-terminal deletion constructs were nonfunctional (**Figure 1E**), neither of these truncation constructs was able to bind FIP200 (**supp Figure 5D**). This is somewhat surprising as we expected the C-terminal deletion mutant to still bind FIP200. We suspect that truncating TNIP1 disturbs its folding and thus its ability to bind FIP200. In order to exclude the possibility that TNIP1 could pull down FIP200 via a secondary interaction with mATG8 binding, we performed CID experiments coupled with immunocytochemistry in a hATG8 6KO background (**supp Figure 5E)**. TNIP1 was able to recruit FIP200 upon RAPALOG treatment, indicating that the binding between TNIP1 and FIP200 is not mediated by hATG8 proteins. The LIR mutant TNIP1, however, was not able to recruit FIP200, confirming that this region is necessary for FIP200 binding (**supp Figure 5F**). To further characterize the interaction between TNIP1 and FIP200, we looked to identify the binding region of TNIP1 within FIP200 (**Figure 5D**). While TNIP1 was able to bind full length and the C-terminal region of FIP200, it was not able to bind the N-terminal part of FIP200 (**Figure 5E**), prompting us to further map the C-terminal region interaction with FIP200. NDP52 was previously shown to interact with the Leucine Zipper domain of FIP200, while the CLAW domain of FIP200 was recently identified as responsible for binding with p62^25^. Consequently, we investigated binding to TNIP1 of these two domains of FIP200. TNIP1 was able to pull-down only the Claw domain of FIP200, indicating that the CLAW domain is the minimal necessary region for binding to TNIP1 (**Figure 5E**).

We next investigated whether this LIR motif was important for TNIP1’s mitophagy function using CID experiments. The LIR mutant TNIP1 was able to trigger mitophagy with an approximately 60% defect compared to WT TNIP1, indicating the importance of the LIR domain in mitophagy (**Figure 6A**). Immunocytochemistry confirmed the FACS data, with the LIR mutant construct recruiting LC3B to the mitochondria to a much lesser extent than WT TNIP1 (**supp Figure 6A, B**). Since we discovered that the LIR mutant and AHD3 deletion mutant TNIP1 were deficient in mitophagy, we combined both and made an AHD3 deletion and LIR mutant construct. This construct completely abolished TNIP1’s ability to induce mitophagy (**Figure 6A**). We conclude that the LIR motif and AHD3 domain of TNIP1 are both involved with distinct roles in mitophagy. We next carried out rescue experiments with the LIR mutant construct in PINK1-Parkin dependent mitophagy. Similarly to the observations with the AHD3 deletion construct, the inhibition of mitophagy was abolished with the LIR mutant construct (**Figure 6B**). Therefore, we conclude that the LIR motif and the AHD3 domain of TNIP1 are both contributing to its role in inhibiting mitophagy.

**Fig. 6.**
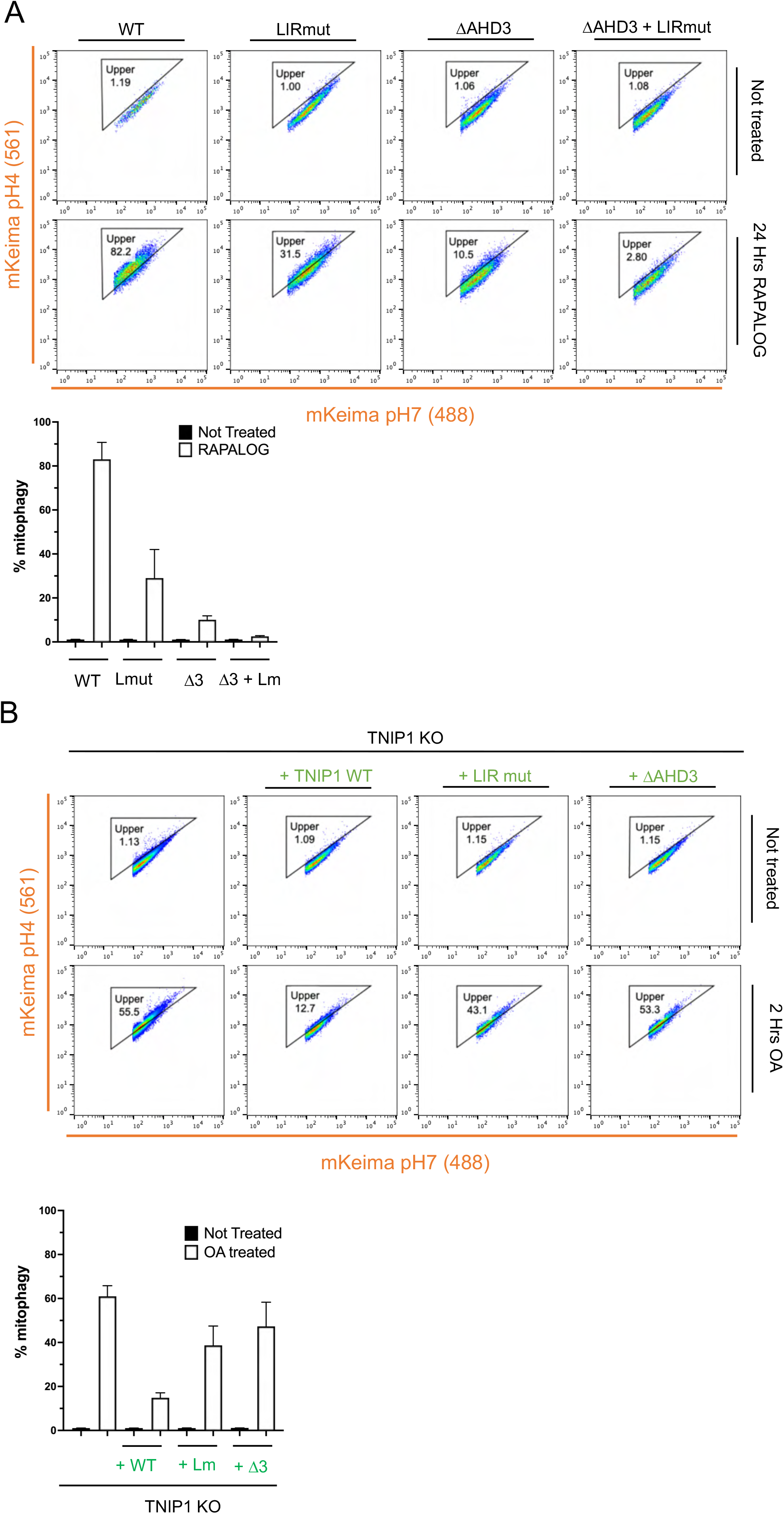
TNIP1 binds FIP200 via its LIR motif. (A) Cells were treated with RAPALOG for 24 Hrs and subjected to FACS acquisition. Quantification of mtKeima ratiometric analysis (561/488) of HeLa cells stably expressing mito-Keima, FRB-FIS1 and FKBP-GFP-TNIP1 WT, LIR mutant, ΔAHD3 or LIR mutant and ΔAHD3 of FACS data from 3 independent replicates. WT: Wild type. Lm: LIR mutant (AEVA), Δ3: ΔAHD3, Δ3+Lm: ΔAHD3 + LIR mutant. (B) Cells were treated with Oligomycin and Antimycin (O/A) for 2 Hrs and subjected to FACS acquisition. Quantification of mtKeima ratiometric analysis (561/488) TNIP1 KO HeLa cells stably expressing mito-Keima, HA-Parkin and FKBP-GFP-TNIP1 wild type, LIR mutant or ΔAHD3 of FACS data from 3 independent replicates. WT: Wild type, Lm: LIR mutant (AEVA), Δ3: ΔAHD3. These cells do not express the FRB-FIS1 construct and thus are not able to perform CID experiments.

## Discussion

Here, we investigated the involvement of the NF-kB pathway-associated protein TNIP1 in autophagy and identify it as a novel inhibitor of mitophagy. TNIP1-mediated mitophagy inhibition relies on binding to both TAX1BP1 (via its AHD3 domain) and FIP200 (via a previously unrecognized LIR motif). Our findings have potentially important implications for our understanding of early events of mitophagy induction, ULK1 complex recycling at the forming autophagosome, and the development of neurodegenerative diseases.

Because TNIP1 decreases the rate of mitophagy, and binds to FIP200, we hypothesize that it acts by sequestering FIP200, and thereby slowing down the mitophagy response. We anticipate this sequestration can happen in two main places during mitophagy (**supp Figure 6C**). (1) TNIP1 binds FIP200 at all times, regulating its availability, preventing its recruitment to autophagy receptors in non-mitophagy conditions. (2) TNIP1 is recruited to the forming autophagosome via TAX1BP1 binding and competes with the autophagy receptors for FIP200 binding, releasing the ULK1 complex from the mitochondria. This hypothesis is supported by the data showing that FIP200 and TNIP1 are not degraded during mitophagy, hence it is most probably removed from the autophagosome before closure, in a mechanism that is still elusive. In this model, the data showing that overexpressing TNIP1 slows down mitophagy is hypothesized to be due to an excess of TNIP1 located to the autophagosome, subsequently releasing the ULK1 complex before it can recruit downstream autophagy components. The findings that ectopic TNIP1 doesn’t inhibit bulk autophagy during starvation is an indication that hypothesis (1), where TNIP1 binds FIP200 at all times, is probably not likely, otherwise an inhibition upon starvation would occur as well. One of the key findings of this report is the identification of a LIR motif in TNIP1. However, similarly to a recent publication describing the FIP200 binding region of SQSTM1/p62 encompassing the LIR motif^25^, it is very likely that a longer sequence than the LIR is responsible for binding to FIP200 and could potentially outcompete LIR binding to mATG8. In this instance, TNIP1 could be driven to the forming autophagosome by binding to the GABARAP proteins, and then compete with the autophagy receptors (i.e., TAX1BP1) for FIP200 binding. The MS data with the ΔAHD3 and ΔAHD4 mutants showing a stronger binding to FIP200 could reflect this binding competition between TNIP1, TAX1BP1 (or OPTN) and FIP200, as it seems that losing the domains responsible for binding to the autophagy receptors enhance FIP200 binding. The tighter binding of ΔAHD3 and ΔAHD4 mutants to FIP200 could also indicate that conformational changes in TNIP1 may regulate mitophagy inhibition. Further work will be necessary to fully understand how TNIP1 is regulated to perform its negative action on mitophagy. Furthermore, future studies on TNIP1 should investigate other types of selective autophagy involved in neurodegenerative diseases such as aggrephagy.

Interestingly, this inhibition of mitophagy mediated by TNIP1 is reminiscent of its role in the negative regulation of the NF-kB pathway or interferon response as well as apoptosis^8,12,20,31^, although for mitophagy it seems that TNIP1 is acting independently of ubiquitin signaling as its UBAN domain is largely not required. To date, very few proteins have been characterized as inhibitors of autophagy. The Deubiquitin enzyme (DUB) USP30 was shown to oppose Parkin by removing ubiquitin molecules from mitochondria^18^. Most other proteins identified negatively regulate bulk autophagy and not selective autophagy. This is exemplified by the negative regulation via the production of PI3P, where several proteins were shown to act. The antiapoptotic protein Bcl-2 was shown to interact and inhibit Beclin-1’s function in autophagy^32^. The PI3P phosphatase Jumpy was shown to control autophagy initiation by acting at the autophagic isolation membrane stage^33^. Autophagy is also inhibited at the LC3 level, with FLIP altering the interaction between ATG3 and LC3^34^. Also acting at the LC3 level, UBA6-BIRC6 was shown to ubiquitinate LC3, regulating it’s availability for autophagy and resulting in an inhibition^35^. Finally, further down the pathway, the Beclin-1 interacting protein Rubicon was shown to negatively regulate autophagosome maturation^36,37^. TNIP1 thus is another addition to these handful described negative regulator of autophagy, albeit TNIP1, like Usp30, seems to be downregulating mitophagy and not bulk autophagy.

Although to our knowledge this is the first time that a negative function of mitophagy is described for TNIP1, it was previously found in proteomics papers as being autophagy substrate^21,38^. TNIP1 was also identified as an autophagy substrate in senescent cells^11^ as well as upon stimulation by Toll-like receptor ligands^39^. It should be noted that we do not see TNIP1 degradation by the autophagy-lysosomal pathway upon mitophagy induction, so its turn-over during autophagy is likely negligible. Furthermore, although we confirmed that A20 was not involved in the negative regulation of mitophagy, A20 was previously shown to interact with ATG16L1, together controlling intestinal homeostasis^40^. However, TNIP1 was not investigated in that study, thus its involvement there is unknown.

Recent findings implicate mitochondrial dysfunction in neurodegenerative disease (ND), and more particularly in Parkinson’s disease (PD) and AD. Although most reports show that a defect in mitophagy is deleterious for ND, and that TNIP1 negatively regulates mitophagy rates, one can hypothesize that loss of TNIP1’s function in mitophagy regulation could be a factor leading to ND such as AD. For these reasons, efforts will be necessary to investigate whether and how TNIP1 could be involved in mitochondrial dysfunction. It will be particularly interesting to see if TNIP1’s locus associated with AD is due to it’s loss of function in mitophagy or any of the other pathways it is involved in. On the other hand, for PD, efforts are being made to activate mitophagy. One strategy seen as amenable to pharmacologic manipulation is to inhibit the proteins that inhibit mitophagy such as USP30. One could consider pharmacologic inhibition of TNIP1 as an alternate approach.

## Acknowledgement

The authors would like to thank the laboratory of Michael Lazarou (Monash Biomedicine Discovery Institute) for sharing the LC3/GABARAP KO cells. We would also like to thank all other members of the Youle lab for sharing cell lines and reagents as well as discussions and advice. We further thank Dr Yan Wang from the National Insitute of Dental and Craniofacial Research Mass Spectrometry Facility (ZIA DE00075) for performing mass spectrometry anlysis. FACS data were acquired using the NINDS facility under the supervision of Dr. Dragan Maric. This work was supported by the Intramural Program of NINDS, NIH.

## Author’s contribution

FLG conceived the study, performed all experiments, analyze and interpreted data. AW performed mass spectrometry and analyzed MS data. CW provided reagents and cell lines as well as intellectual contribution. FLG and RY wrote the manuscript with input from all authors. RY supervised the project and acquired funds.

## Declaration of Interest

The authors declare no competing interests.

## Figure Legends

**Supplementary Fig. 1.**
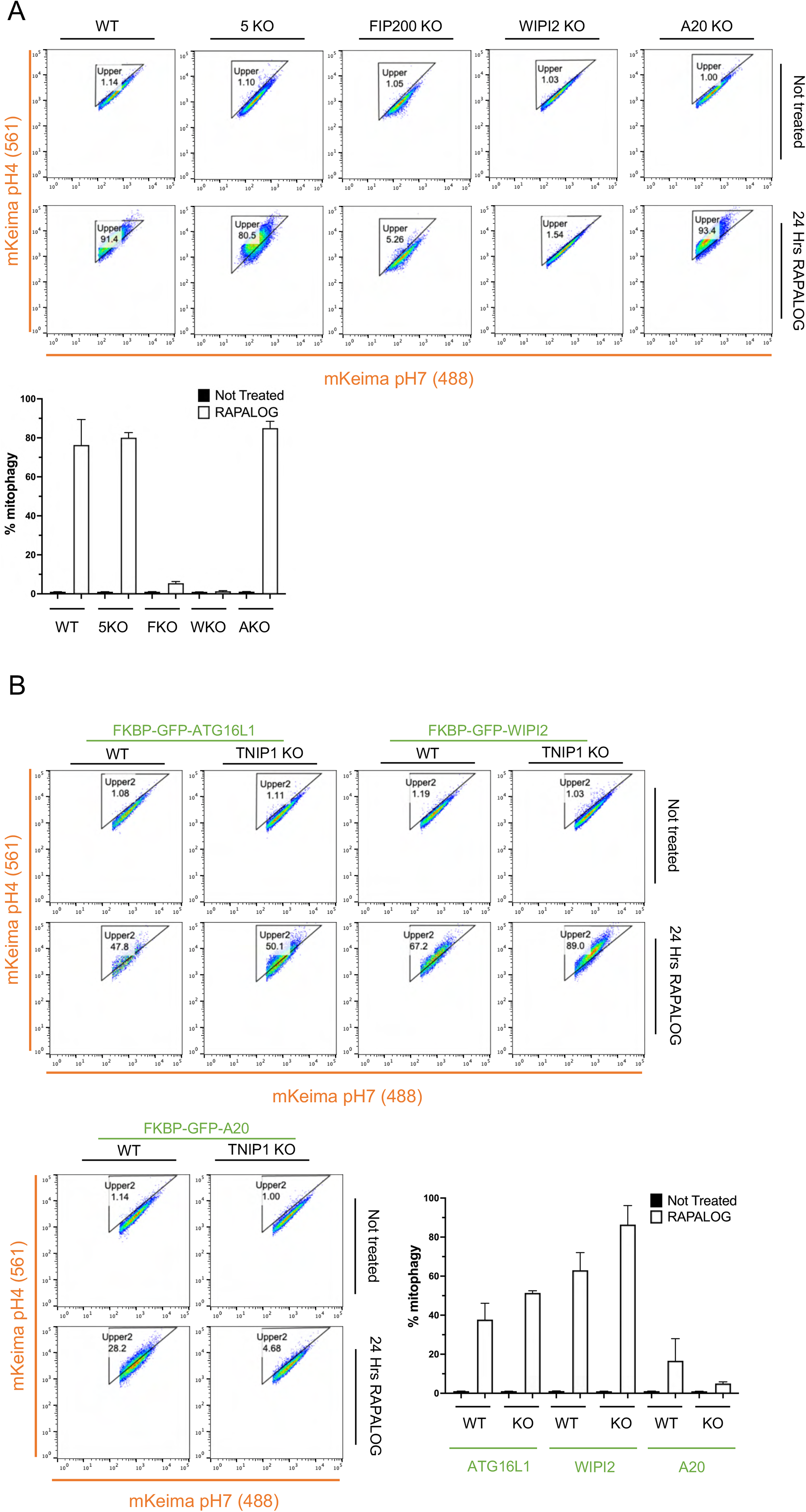
Mitophagy by ectopic localization of TNIP1 to the mitochondria is dependent of the autophagy machinery. (A) Top, cells were treated with RAPALOG for 24 Hrs and subjected to FACS acquisition. FACS plot of mtKeima ratio (561/488) of HeLa cells stably expressing mito-Keima, FRB-FIS1 and FKBP-GFP-TNIP1 in wild type, 5KO (SQSTM1, NBR1, NDP52, TAX1BP1 and TAX1BP1 KO cells), FIP200 KO, WIPI2KO or A20 KO cells. Bottom, quantification of mtKeima ratiometric analysis of FACS data from 3 independent replicates. WT: Wild type. K KO: FIP200 KO, W KO: WIPI2KO and A KO: A20 KO. (B) Top, cells were treated with RAPALOG for 24 Hrs and subjected to FACS acquisition. FACS plot of mtKeima ratio (561/488) of HeLa cells stably expressing mito-Keima, FRB-FIS1 and FKBP-GFP-ATG16L1 or FKBP-GFP-WIPI2 or FKBP-GFP-A20 in wild type or TNIP1 KO cells. Bottom, quantification of mtKeima ratiometric analysis of FACS data from 3 independent replicates. WT: Wild type. KO: TNIP1KO.

**Supplementary Fig. 2.**
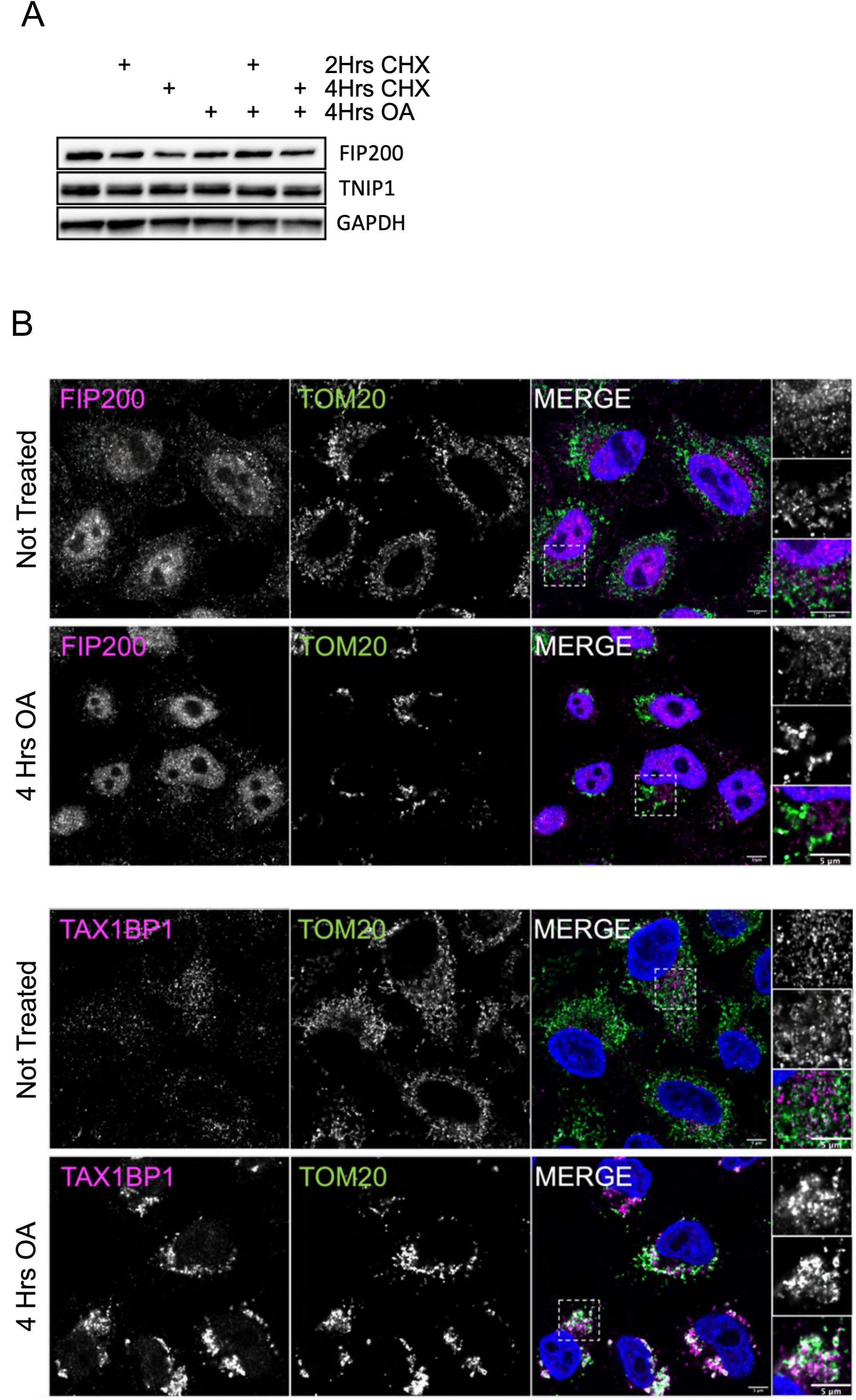
(A) HeLa cells stably expressing HA-Parkin were treated for 4 Hrs with Oligomycin and Antimycin (O/A) and/or 2 Hrs and 4 Hrs of cycloheximide (CHX) and subjected to immunoblot analysis. (B) HeLa cells stably expressing HA-Parkin were treated for 4 Hrs with Oligomycin and Antimycin (O/A) and stained for the mitochondrial protein TOM20 and endogenous FIP200 and TAX1BP1 before immunofluorescence acquisition on a confocal microscope. Airyscan representative images. Scale bar: 5μm.

**Supplementary Fig. 3.**
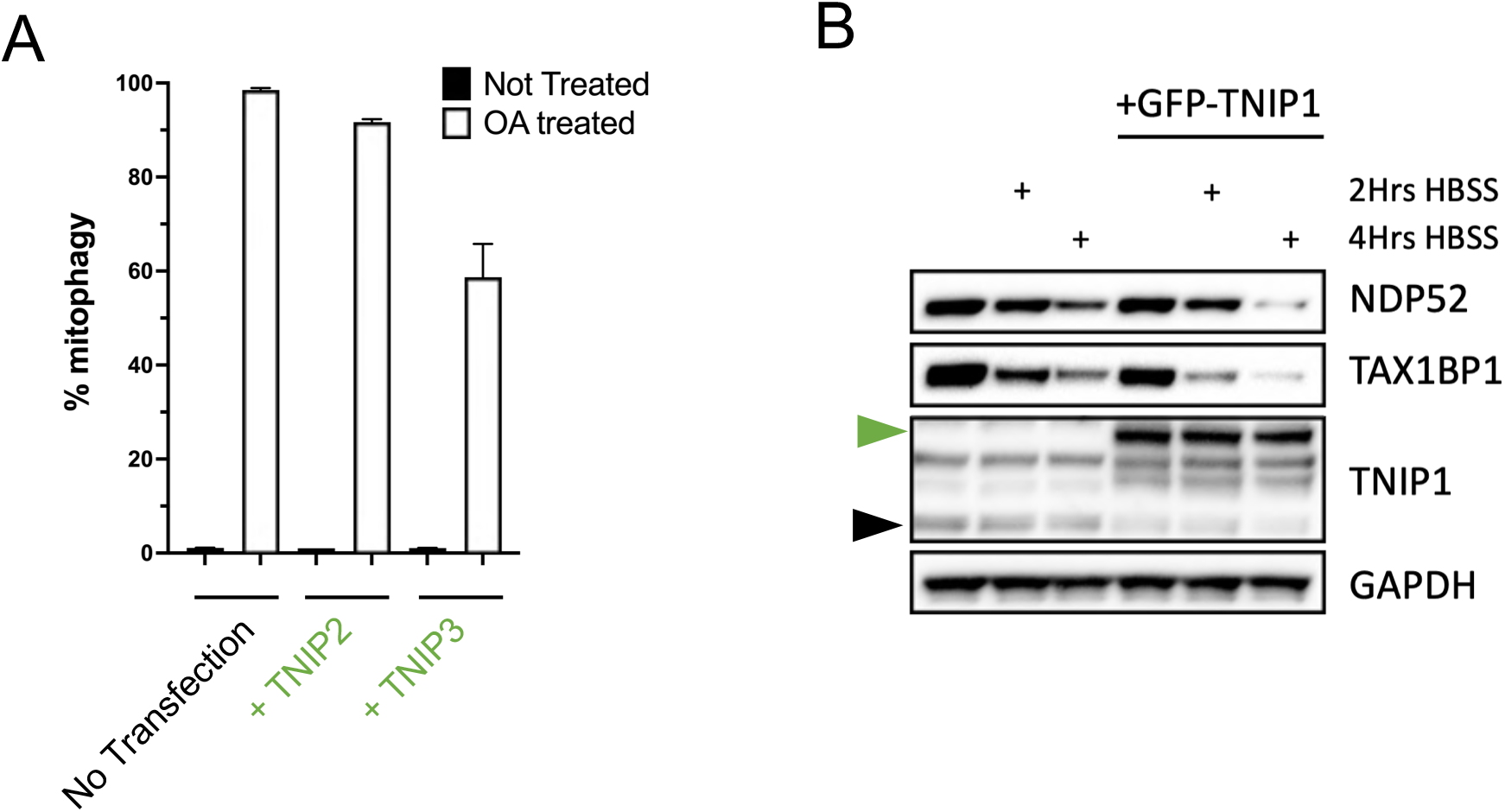
(A) Cells were treated with Oligomycin and Antimycin (O/A) for 6 Hrs and subjected to FACS acquisition. Quantification of mtKeima ratiometric analysis (561/488) of HeLa cells stably expressing mito-Keima, HA-Parkin and FKBP-GFP-TNIP2 or FKBP-GFP-TNIP3 of FACS data from 3 independent replicates. NT: Not transfected wild type. T2: FKBP-GFP-TNIP2 transfected cells. T3: FKBP-GFP-TNIP3 transfected cells. (B) HeLa cells stably expressing FKBP-GFP-TNIP1 WT were starved for 2 Hrs, or 4 Hrs by incubation with HBSS and subjected to immunoblot analysis.

**Supplementary Fig. 4.**
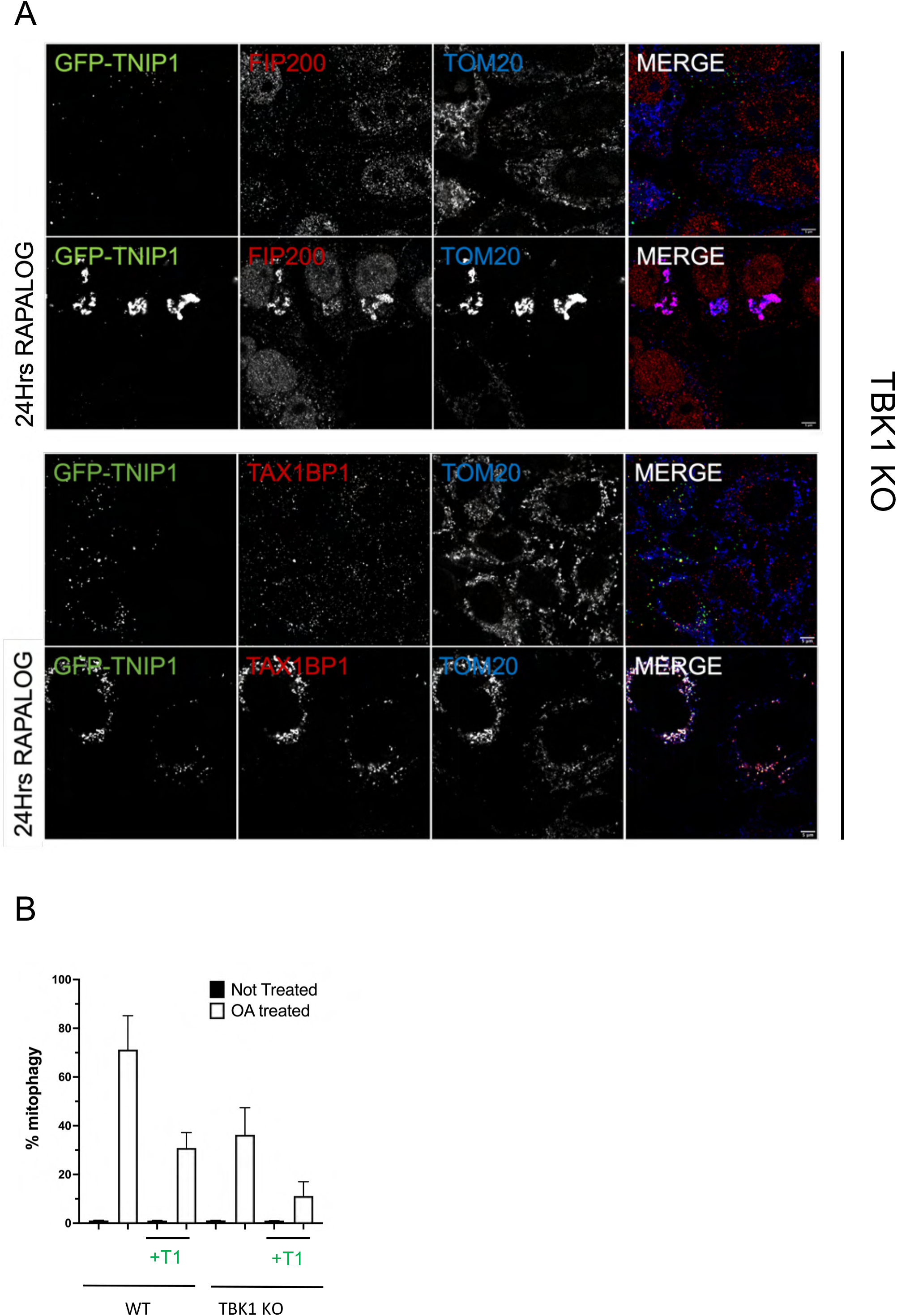
TBK1 is not involved in TNIP1-mediated inhibition of mitophagy. (A) TBK1 KO HeLa cells stably expressing FRB-FIS1 and FKBP-GFP-TNIP1 were treated for 24 Hrs with RAPALOG and stained for the mitochondrial protein TOM20 and endogenous FIP200 or TAX1BP1 before immunofluorescence acquisition on a confocal microscope. Airyscan representative images. Scale bar: 5μm. (B) Cells were treated with Oligomycin and Antimycin (O/A) for 6 Hrs and subjected to FACS acquisition. Quantification of mtKeima ratiometric analysis (561/488) of WT or TBK1 KO HeLa cells stably expressing mito-Keima, HA-Parkin and FKBP-GFP-TNIP1 of FACS data from 3 independent replicates. WT: Wild type. + T1: + FKBP-GFP-TNIP1. These cells do not express the FRB-FIS1 construct and thus are not able to perform CID experiments.

**Supplementary Fig. 5.**
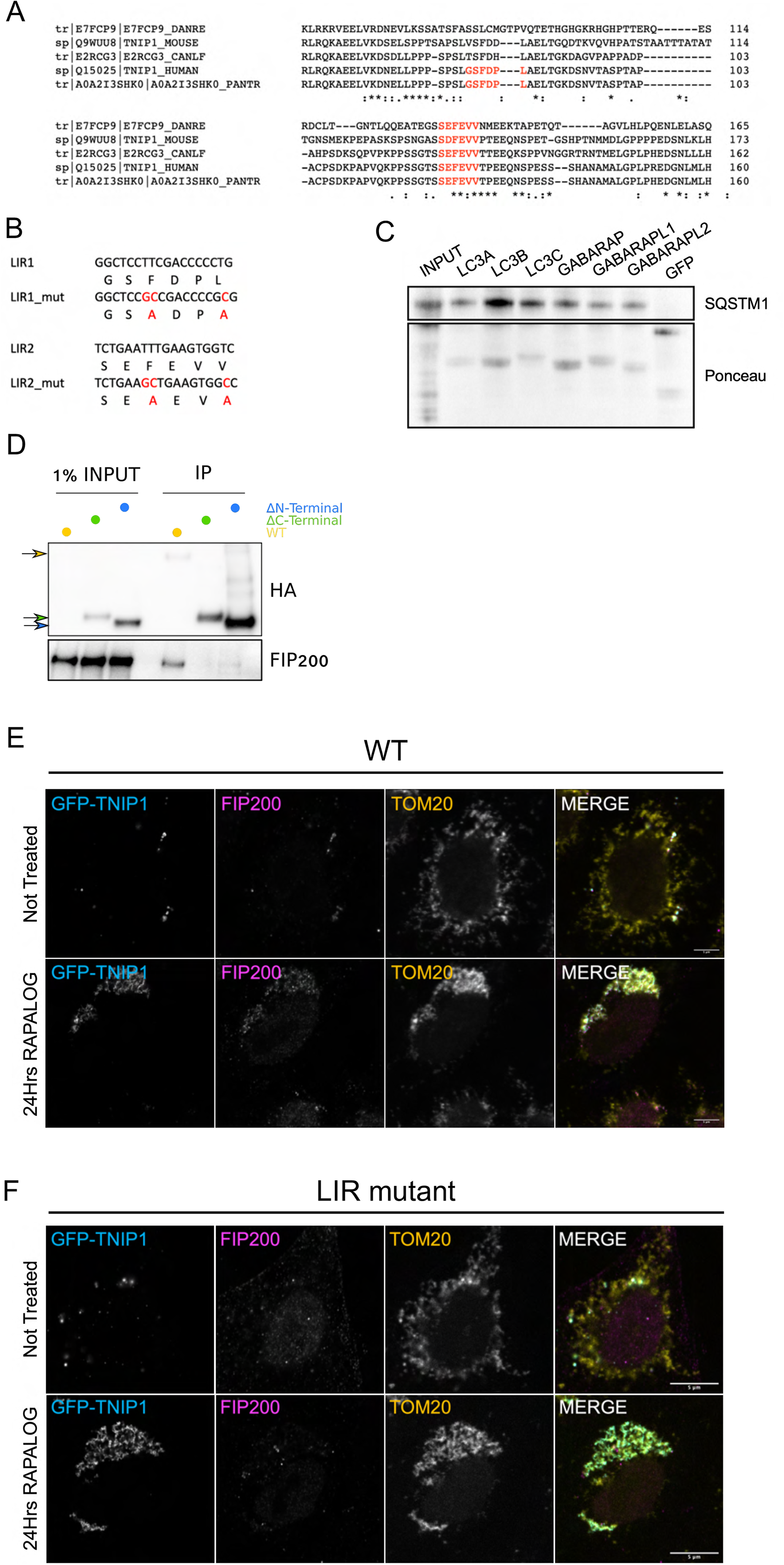
(A) Alignment sequence analysis of Human, Zebrafish (DANRE), Mouse, Dog (CANLF) and Chimpanzee (PANTR). Highlighted in red are the identified putative LIR motifs. (B) sequence showing the nucleic acids mutated replaced to obtain alanines and alter the LIR motif. (C) HeLa cells stably expressing HA-SQSTM1 were lysed and lysate was incubated with purified recombinant GST tagged mATG8 proteins of GFP. Precipitates were subjected to immunoblot analysis. (D) HEK293T cells stably expressing HA-TNIP1 full length or the ΔC-terminal and ΔN-terminal constructs were immunoprecipitated using magnetic HA beads and subjected to immunoblot analysis. (E) HeLa cells stably expressing FRB-FIS1 and FKBP-GFP-TNIP1 were treated for 24 Hrs with RAPALOG and stained for the mitochondrial protein TOM20 and endogenous FIP200 before immunofluorescence acquisition on a confocal microscope. Airyscan representative images. Scale bar: 5μm. (F) HeLa cells stably expressing FRB-FIS1 and FKBP-GFP-TNIP1 LIR mutant were treated for 24 Hrs with RAPALOG and stained for the mitochondrial protein TOM20 and endogenous FIP200 before immunofluorescence acquisition on a confocal microscope. Airyscan representative images. Scale bar: 5μm.

**Supplementary Fig. 6.**
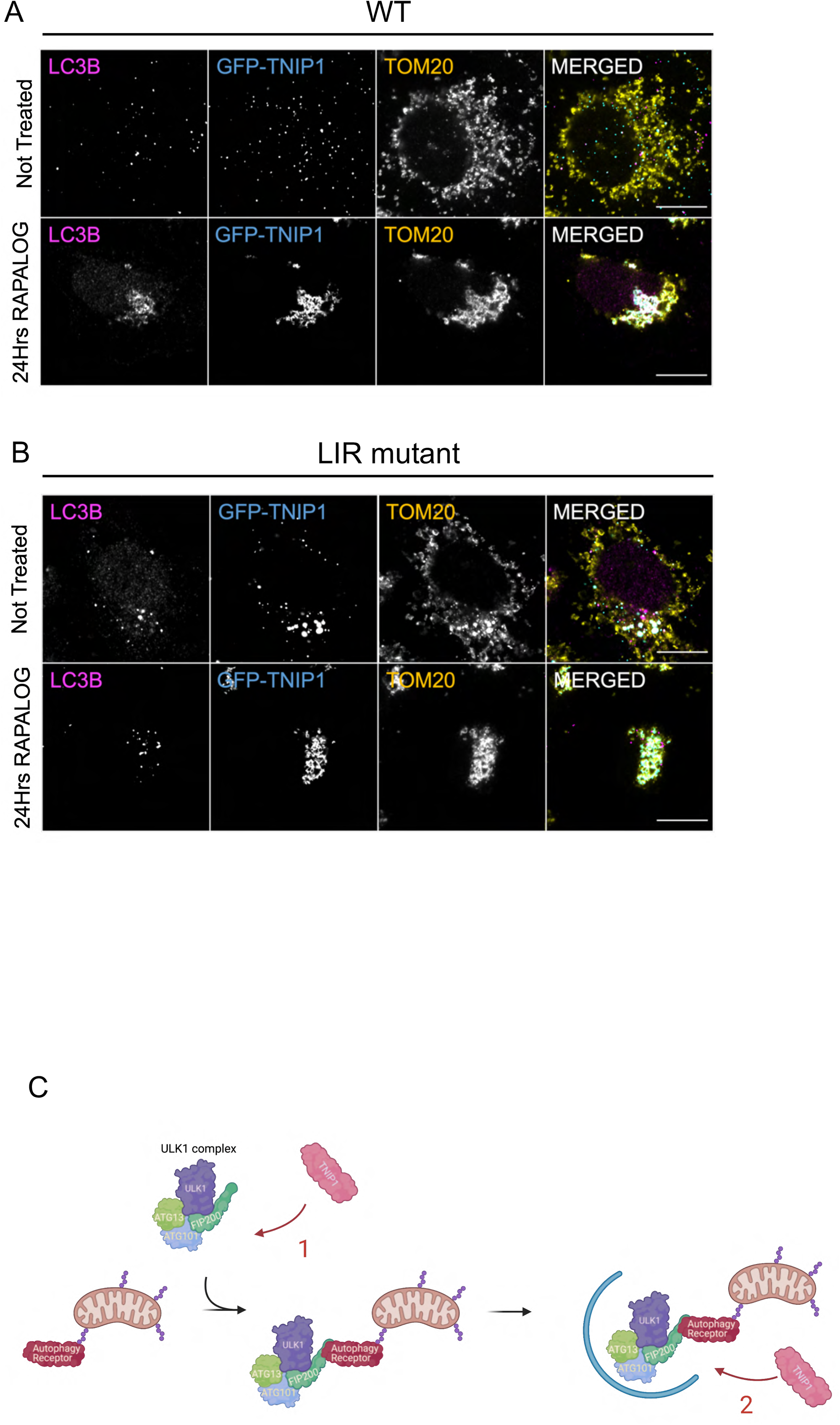
(A) HeLa cells stably expressing FRB-FIS1 and FKBP-GFP-TNIP1 were treated for 24 Hrs with RAPALOG and stained for the mitochondrial protein TOM20 and endogenous LC3B before immunofluorescence acquisition on a confocal microscope. Airyscan representative images. Scale bar: 5μm. (B) HeLa cells stably expressing FRB-FIS1 and FKBP-GFP-TNIP1 LIR mutant were treated for 24 Hrs with RAPALOG and stained for the mitochondrial protein TOM20 and endogenous LC3B before immunofluorescence acquisition on a confocal microscope. Airyscan representative images. Scale bar: 5μm. (C) Schematic representation of the two possible mode of inhibition of mitophagy by TNIP1, made with the help of BioRender. 1, TNIP1 sequesters FIP200 and prevents its recruitment to the mitochondria. 2, TNIP1 is recruited to the mitochondria and displaces the ULK1 complex via comepition binding between the autophagy receptor and FIP200.

## Material and Methods

### Reagents or Resource

#### Antibodies

A20 Bethyl A302-633A

Actin CST 4967S

ATG16L1 CST 8089

COXII abcam ab110258

FIP200 CST 12436

FIP200 Proteintech 17250-1-AP

GAPDH G9545

GFP Invitrogen A11122

HA CST 3724s

HA biolegend 901513

HA MMS-101P

LC3B CST 2775s

LC3B PM036

MFN2 House made

NDP52 CST 60732S

OPTN proteintech10837-I-AP

SQSTM1 abnova H00008878-M01

TAX1BP1 CST 5105S

TAX1BP1 HPA024432

TNIP1 Bethyl A304-508

TNIP1 proteintech 15104-1-AP

TOM20 SC-11415

TOM20 SC-17764

WIPI2 abcam ab105459

Alexa Fluor 488 goat anti-rabbit IgG Thermofisher A11008

Alexa Fluor 594 goat anti-rabbit IgG Thermofisher A11012

Alexa Fluor 405 goat anti-mouse IgG Thermofisher A31553

Alexa Fluor 488 goat anti-mouse IgG Thermofisher A21202

Mouse secondary

Rabbit secondary

#### Chemicals, peptides and recombinant proteins

A/C heterodimerizer (Rapalog)

antimycin A

Bafilomycin

DAPI

Oligomycin

Cycloheximide

Polybrene

Q-VD

Sigma FBS

DMEM

Sodium pyruvate

GlutaMax

Opti-MEM

0.05% Trypsin-EDTA

BamBanker

DTT

HBSS with calcium and magnesium

paraformaldehyde 16% solution, EM grade

polyethylenimine, linear, MW 25000, transfection grade, (PEI 25K)

XtremeGene

Protease inhibitor cocktail

Puromycin

Triton X-100

Tween-20

3xHA peptide

MS grade trypsin

MMTS

TEAB

#### Commercial assays

Amersham ECL Prime WB detection reagent

BCA

SuperSignal West Femto

NEBuilderHiFi DNA assembly master mix

Q5 Hot Start High-Fidelity DNA Polymerase

DNA Ligation kit, Mighty Mix

Gateway BP clonase

Gateway LR clonase

LDS sample buffer (4x)

Chromotek GFP-TRAP magnetic beads

Thermofisher HA Beads

S-Trap column

PlasmoTest

Pre-treated 1.5mm coverslips GG-18-1.5-pre (neuVitro)

#### Deposited data

MS data

#### Experimental models: Cell lines

HeLa

293T

HeLa FIP200 KO

HeLa WIPI2 KO

HeLa A20 KO

HeLa TNIP1 KO

HeLa autophagy receptor 5KO

HeLa TAX1BP1 KO

HeLa ATG8 6KO

HeLa TBK1 KO

#### Oligonucleotide

TNIP1 sgRNA GACCCCCTGGCTGAGCTCAC

TAX1BP1 sgRNA AAUUGUGUACUAGCAUUCCA

A20 sgRNA GGCTGAACAAGTCCTTCCTC

#### Recombinant DNA

pENTR223-TNIP1

pENTR223-TNIP1 LIR1mut

pENTR223-TNIP1 LIR2mut

pHAGE-FLAG-HA TNIP1 WT

pHAGE-FLAG-HA TNIP1 LIR1 mutant

pHAGE-FLAG-HA TNIP1 LIR2 mutant

pHAGE-FLAG-HA TNIP1 ΔAHD1

pHAGE-FLAG-HA TNIP1 ΔAHD3

pHAGE-FLAG-HA TNIP1 ΔAHD4

pHAGE-FLAG-HA TNIP1 ΔCterminal

pHAGE-FLAG-HA TNIP1 ΔNterminal

pHAGE-FLAG-HA TAX1BP1 WT

pHAGE-FLAG-HA TAX1BP1 CC

pHAGE-FLAG-HA TAX1BP1 ΔSKITCH

pHAGE-FLAG-HA TAX1BP1 ΔZF

pHAGE-FLAG-HA SQSTM1

pHAGE-FLAG-HA FIP200 ΔCterminal

pHAGE-FLAG-HA FIP200 ΔNterminal

pHAGE-FLAG-HA FIP200 LZ

pHAGE-FLAG-HA FIP200 CLAW

pHAGE-FKBP-GFP-TNIP1 WT

pHAGE-FKBP-GFP-TNIP1 D472N

pHAGE-FKBP-GFP-TNIP1 LIR2 mutant

pHAGE-FKBP-GFP-TNIP2

pHAGE-FKBP-GFP-TNIP3

pHAGE-FKBP-GFP-TNIP1 ΔAHD1

pHAGE-FKBP-GFP-TNIP1 ΔAHD3

pHAGE-FKBP-GFP-TNIP1 ΔAHD4

pHAGE-FKBP-GFP-TNIP1 ΔAHD3+LIR2 mutant

pHAGE-FKBP-GFP-TNIP1 ΔCterminal

pHAGE-FKBP-GFP-TNIP1 ΔNterminal

pHAGE-FKBP-GFP-ATG16L1

pHAGE-FKBP-GFP-WIPI2

pHAGE-FKBP-GFP-A20

pME18s-3xHA-FIP200

pSpCas9-P2A-Puro

mtKeima-P2A-HA-Parkin

#### Software and algorithm

Fiji

Prism

FlowJo

SnapGene

Image Lab

BioRender

### Experimental Model and Subject Detail

#### Cell line

HEK293T and HeLa cells were purchased from ATCC. HEK293T and HeLa cells were cultured in Dulbecco’s modified eagle medium (DMEM) with 10% (v/v) Fetal Bovine Serum (FBS) (Sigma), 1 mM Sodium Pyruvate, 2 mM GlutaMAX. All media and supplements were from Thermo Fisher. All cells were tested for mycoplasma contamination every two weeks with PlasmoTest kit (InvivoGen). Reagents used for transfections were, X-tremeGENE 9 (Roche) for sgRNA transfection, or Polyethylenimine (PEI) (PolySciences) for all other transfections. The full list of antibodies and reagents are found in Key Resources Table.

### Method Detail

#### Knockout line generation using CRISPR/Cas9 gene editing

CRISPR gRNAs were generated to target exon 3 of TNIP1, exon 2 of TNFAIP3 and exon 3 of TAX1BP1. gRNAs were cloned into pSpCas9(BB)-2A-Puro (PX459) V2.0 (Addgene plasmid #62988). To Make KO, HeLa cells were transfected with the gRNA plasmid and treated with 1 ug/ml puromycin for 2 days to enrich transfected cells, which were then diluted and placed into 96-well plates for single colonies. Primer set (ggtggaccagcatggagttt and accagggagcttccaactca) were used for PCR screening of TNIP1 KO clones. Primer set (tcagtacccactctctgccttc and ctccaagcctcaatgtgctct) were used for PCR screening of TNFAIP3 KO clones. Primer set (ttatccttgagaaattggatagca and tagtacctaaaaagaaacccactcttc) were used for PCR screening of TAX1BP1 KO clones.

#### Cloning and stable cell line generation

For lentiviral constructs, inserts were either amplified by PCR and cloned into pHAGE vector, respectively by Gibson assembly (New England Labs) or Gateway cloning (Thermo Fisher). Deletion mutants were generated using Gibson Cloning. Point mutants were generated by site directed mutagenesis. All constructs used or generated in this study were validated by Sanger sequencing and complete plasmid sequences and maps are available upon request.

Stable expression of lentiviral constructs in HeLa or HEK293T cells were achieved as follows: lentiviruses were packaged in HEK293T cells by transfecting constructs together with appropriate helper plasmids and PEI. The next day, media was exchanged with fresh media. Viruses were harvested 48 hrs and 72 hrs after transfection and transduced in HeLa or HEK293T cells with 8 μg/mL polybrene (Sigma). Cells were then directly used in experiments or optimized for expression by FACS.

#### Immunoblot Analyses

Cells seeded into 6-well plates were washed with phosphate buffered saline (PBS) and lysed with RIPA buffer. The protein concentration was measured using a BCA kit. Samples were boiled at 99°C for 10 min. 20-50 μg of protein lysate of each sample was loaded and separated on 4-12% Bis-Tris gels (Thermo Fisher or GenScript) according to manufacturer’s protocol. Gels were transferred to polyvinyl difluoride membranes and immunostained using specific antibodies. For mitophagy measurements by immunoblotting, cells were treated with 10 μM Oligomycin (Calbiochem), 10 μM Antimycin A (Sigma) and 10 μM QVD (ApexBio) in growth medium at different timepoints indicated in figure legends, prior to western blot analysis. For starvation-induced autophagy, by immunoblotting, cells were washed 2 times with PBS and incubated for the indicated time with HBSS containing calcium and magnesium.

#### Recombinant protein production and protein purification

mATG8 proteins (LC3A, LC3B, LC3C, GABARAP, GABARAPL1, GABARAPL2) and GFP were first cloned into pENTR vector using GATEWAY cloning. They were cloned into pDEST60 GST vector using LR clonase. GST expression vector were expressed into DL21 (DE3) bacteria. 50ml LB broth was inoculated with 2ml pre-culture of the GST constructs. Bacteria were incubated for ∼1 Hr at 37°C with agitation until an optical density of ∼0.6 was reach. 400μM IPTG was used to induce protein production for 4 Hrs. Bacteria were collected and the pellet was lysed with lysis buffer (20mM TRIS-HCl pH7.5, 10mM EDTA, 5mM EGTA, 150mM NaCl, 0.5% NP40, 1% Triton X-100, Benzonase, 1mM DTT, protease inhibitor, 2mg/ml lysozyme). The lysate was shockfreezed with liquid nitrogen before thawing and sonication. The samples were centrifuged and cleared lysate was incubated with 100μl of slurry GST beads, followed by an overnight incubation at 4°C on a rotating shaker. Beads were subsequently washed (20mM TRIS-HCl pH 7.5, 10mM EDTA, 5mM EGTA, 150mM NaCl, 1mM DTT) before exchanging the buffer for the storage buffer (20mM TRIS-HCl pH 7.5, 10mM EDTA, 5mM EGTA, 150mM NaCl, 1mM DTT, 5% glycerol, proteinase inhibitor).

#### Immunoprecipitation, GST-pull down, GFP-TRAP and HA beads precipitation

For GFP-TRAP (Chromotek), HA-beads (Pierce), precipitation experiments, HEK293T cells seeded in 10cm plates were co-transfected with specific constructs to overexpress proteins of interest for 24 to 48 h. IPs were performed following manufacturer’s instructions. Briefly, for HA-IP, cells were then lysed using ice cold lysis buffer (25 mM Tris-HCl pH 7.4, 150 mM NaCl, 1% NP-40, 1 mM EDTA, 5% glycerol) supplemented with EDTA-free cOmplete protease inhibitor (Roche). Samples were incubated on ice with intermittent agitation by pipetting for 10 min. Beads were equilibrated using bead resuspension buffer (TBS containing 0.05% Tween-20 Detergent (TBS-T)). Protein lysates were precleared by centrifugation at 4°C for 10 min at 15,000 g. 1% INPUT was collected and clarified lysate was incubated with equilibrated beads for 1-2 Hrs at 4°C. Beads were then washed with ice cold TBST Wash buffer 3 to 5 times. Bound proteins were eluted with 2x LDS lysis buffer (Thermo Fisher) in boiling conditions.

For GFP-Trap, cells were then lysed using ice cold lysis buffer (10 mM Tris/Cl pH 7.5, 150 mM NaCl, 0.5 mM EDTA, 0.5 % NonidetTM P40 Substitute) supplemented with EDTA-free cOmplete protease inhibitor (Roche). Samples were incubated on ice with intermittent agitation by pipetting for 30 min. Beads were equilibrated using bead dilution buffer (10 mM Tris/Cl pH 7.5, 150 mM NaCl, 0.5 mM EDTA). Protein lysates were precleared by centrifugation at 4°C for 10 min at 15,000 g. Clarified lysates were diluted using dilution buffer, 1% INPUT was collected and lysate was incubated with equilibrated beads for 1-2 Hrs at 4°C. Beads were then washed with ice cold Wash buffer (10 mM Tris/Cl pH 7.5, 150 mM NaCl, 0.05 % NonidetTM P40 Substitute, 0.5 mM EDTA) 3 to 5 times. Bound proteins were eluted with 2x LDS lysis buffer (Thermo Fisher) in boiling conditions.

For GST pull down, cells were lysed (50mM TRIS-HCl 7.5, 150mM NaCl, 0.5% NP40, protease inhibitor). Samples were incubated at 4°C with constant agitation for 30 min. Protein lysates were precleared by centrifugation at 4°C for 10 min at 15,000 g. 1% INPUT was collected and clarified lysate was incubated with equilibrated beads overnight at 4°C with agitation. Beads were washed 5 times with wash buffer (20mM TRIS-HCl pH 7.5, 10mM EDTA, 5mM EGTA, 150mM NaCl, 1mM DTT). Bound proteins were eluted with 2x LDS lysis buffer (Thermo Fisher) in boiling conditions.

#### Immunofluorescence microscopy

Cells seeded in 12 well plates with a 1.5mm pre-treated coverslip (neuVitro) were treated as indicated in figure legends. After treatment, cells were rinsed in PBS and fixed with 4% paraformaldehyde at RT for 10 min. Cells were washed with PBS. For immunostaining, cells were then permeabilzed and blocked with 0.1% Triton X-100, 3% goat serum in PBS for 15 min at RT. After, cells were incubated with 0.1% Triton X-100, 3% goat serum in PBS supplemented with antibodies (1:1000) 1-2 Hrs at room temperature. Cells were then washed with PBST (PBS + 0.1% Triton X-100) 3 times and incubated with Alexa 505, 488 or 594-conjugated secondary antibodies (Thermo Fisher). For cells expressing fluorescent tagged proteins, cells were seeded as above. After treatments, cells were fixed as above and washed 3 times with PBS prior to image analysis. Images were taken using 63x oil, DIC objective on an LSM 880 Airyscan microscope (Zeiss).

#### Mass spectrometry

After samples were bound to magnetic HA-beads, samples were eluted with elution buffer (2mg/ml HA peptide in TBS) according to manufacturer’s instructions, followed by TCA precipitation. Briefly, 100% TCA was added to eluted peptides (f.c. 20% TCA), vortexed and incubated on ice for 30min before centrifugation. Precipitated peptides were washed with 10% ice cold TCA, followed by 4 washes with ice cold acetone. The tubes were left to air dry before trypsin digestion. Briefly, peptides were re-suspended with solubilizing buffer (5% SDS, 8M urea, 50mM TEAB, pH7.6). DTT was added for f.c. 5mM before incubating at 50°C for 30min. MMTS was added for 45min before adding an aqueous phosphoric acid solution. Meanwhile, S-Trap columns (Protifi) were conditioned with binding buffer (90% methanol, 100mM TEAB, pH7.1). 1μg of trypsin was added to the acidified protein mixture and immediately added into the column containing binding buffer. Centrifugation and washing steps were performed before adding digestion buffer (Trypsin in 50mM TEAB) and incubation at 37°C overnight. Rehydrate the column with 50mM TEAB before elution using 0.2% formic acid and another elution using 50% acetonitrile containing 0.2% formic acid.

Tryptic digests were analyzed using an orbitrap Fusion Lumos tribrid mass spectrometer interfaced to an UltiMate3000 RSLC nano HPLC system (Thermo Scientific) using data dependent acquisition. Initial protein identification was carried out using Proteome Discoverer (V2.4) software (Thermo Scientific). Search results from Proteome Discoverer were incorporated into Scaffold4 for relative quantification using spectral counting. Samples were compared to ∼100 reference IPs using a Java script programmed according to the CompPASS software suite as previously described^23,24^ and also compared to the CRAPome^41^. For determination of the TNIP1 interaction network, thresholds for high confidence interaction partners (HCIPs) were top 5% of interactors with highest Z-score and lowest abundance in the CRAPome. Cytoscape was used to visualize the TNIP1 interaction network. To compare WT TNIP1 with TNIP1ΔAHD3 and TNIP1ΔAHD4, total spectral counts for each interactor were first normalized to 1000 and then expressed relative to WT/controls in fold-change (log_2_) and plotted as heatmap using GraphPad.

#### Mitophagy assay with mito-mKeima via fluorescence activated cytometer

Stable cell lines were generated to express mito-mKeima, HA-Parkin, FRB-Fis1 and FKBP-GFP-tagged target genes with lentivirus system. 100K cells were seeded in 12-well plates and treated with Rapalog for 24h or by OA at various time points before FACS analysis. FACS analysis were performed as previously described^22,42^.

#### Quantification and Statistical Analysis

All statistical analyses for FACS were calculated in GraphPad Prism 9. Error bars are expressed as mean ± standard deviation.

## Notes

### Competing Interest Statement

The authors have declared no competing interest.

